# Manipulating neural dynamics to tune motion detection

**DOI:** 10.1101/2021.11.02.466844

**Authors:** Aneysis D. Gonzalez-Suarez, Jacob A. Zavatone-Veth, Juyue Chen, Catherine A. Matulis, Bara A. Badwan, Damon A. Clark

## Abstract

Neurons integrate excitatory and inhibitory signals to produce their outputs, but the role of input timing in this integration remains poorly understood. Motion detection is a paradigmatic example of this integration, since theories of motion detection rely on different delays in visual signals. These delays allow circuits to compare scenes at different times to calculate the direction and speed of motion. It remains untested how response dynamics of individual cell types drive motion detection and velocity sensitivity. Here, we sped up or slowed down specific neuron types in *Drosophila*’s motion detection circuit by manipulating ion channel expression. Altering the dynamics of individual neurons upstream of motion detectors changed their integrating properties and increased their sensitivity to fast or slow visual motion, exposing distinct roles for dynamics in tuning directional signals. A circuit model constrained by data and anatomy reproduced the observed tuning changes. Together, these results reveal how excitatory and inhibitory dynamics jointly tune a canonical circuit computation.

## Introduction

When a neuron integrates synaptic inputs, the dynamics of those inputs are critical to the neuron’s output response. However, the role of neural input dynamics in basic computations remains poorly understood, in part because of difficulties in manipulating neural response dynamics. Previous studies have predominantly manipulated neural dynamics by using temperature and pharmacology (Arenz et al., 2017; Banerjee et al., 2021; Long and Fee, 2008; Suver et al., 2012; Tang et al., 2010), but these methods affect entire circuits, making it difficult to investigate how dynamics of individual excitatory and inhibitory input neurons drive computation. In this study, we use the powerful genetic tools in *Drosophila* to manipulate dynamics of *individual* excitatory and inhibitory visual neuron types to examine how these dynamics tune downstream computations.

Circuits that detect visual motion offer a robust testbed for understanding how excitatory and inhibitory input dynamics contribute to the computations of downstream neurons. To detect motion, neurons must integrate visual information over both space and time. Indeed, theories of visual motion detection require adjacent visual signals to be processed with different delays to generate direction-selective responses (Adelson and Bergen, 1985; Barlow and Levick, 1965; Hassenstein and Reichardt, 1956) (Fig. 1A). In both vertebrates and invertebrates, these different delays are thought to be implemented through the response dynamics of neurons upstream of motion-detecting cells (Arenz et al., 2017; Kim et al., 2014). However, it remains untested how the dynamics of upstream excitatory and inhibitory neurons drive downstream motion signals. Motion computation is a compelling framework for investigating this question because motion signals are highly interpretable in their selectivity for direction and speed of motion.

**Figure 1.**
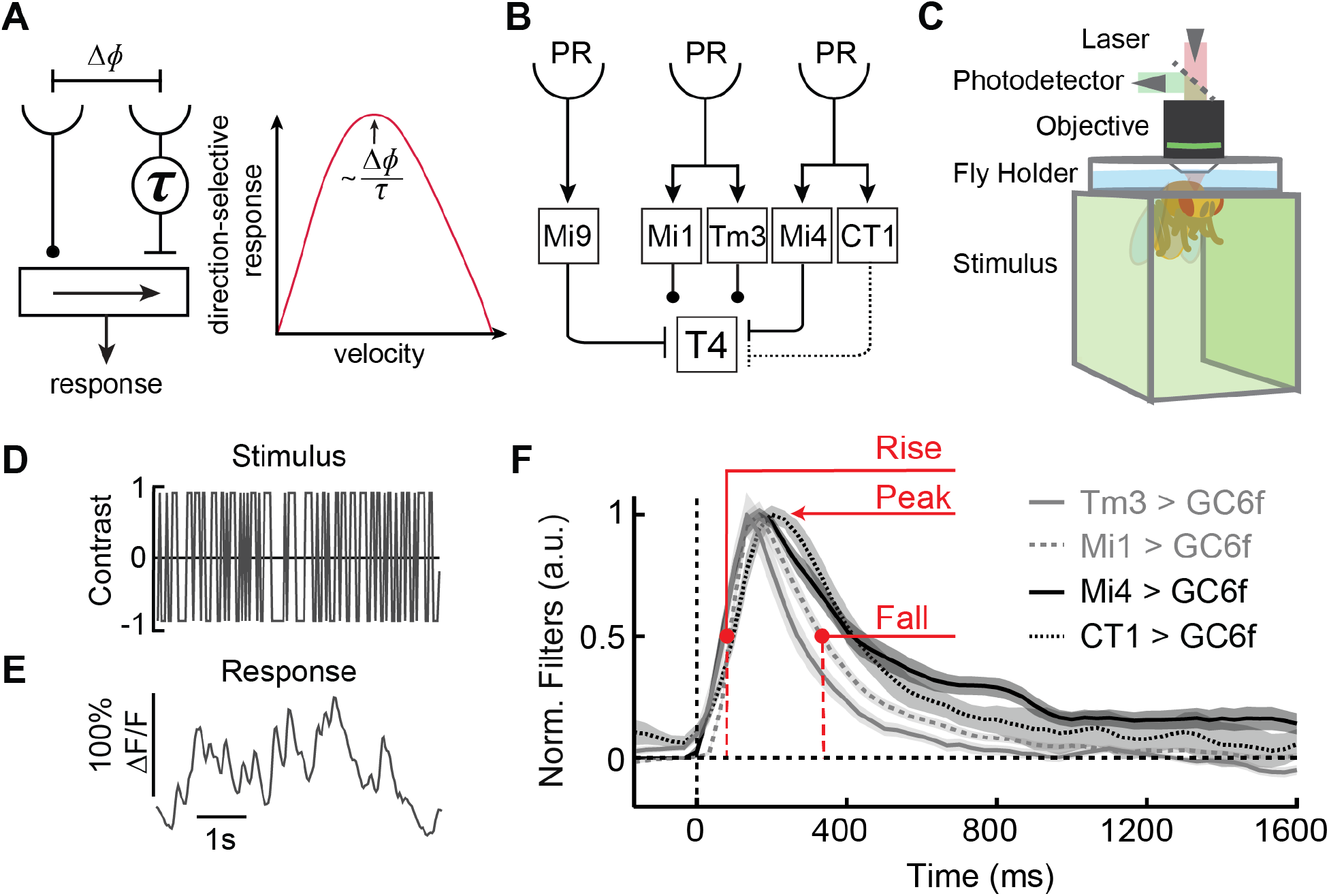
Medulla neurons exhibit heterogeneous filter dynamics. **(A)** A model of direction-selective motion detection has two inputs separated by an angle *Δϕ*, one of which is delayed by a time *τ* Nonlinear combination of the two signals results in a direction-selective response. The model’s direction-selective response is maximal at a velocity that scales with the sensor separation (*Δϕ*) divided by the temporal delay (*τ*). **(B)** Circuit diagram highlighting neurons with strong anatomical connections to the direction-selective cell T4 (Takemura et al., 2017). Solid lines highlight connections that have been established functionally (Strother et al., 2017), while the dashed line refers to an anatomical connection without established function. **(C)** Two-photon imaging was performed in head-fixed flies viewing stimuli presented on panoramic screens. **(D)** A stochastic, binary, high-contrast stimulus was presented to flies to facilitate estimating neural linear filtering properties. **(E)** Example response trace to the stimulus of an Mi1 neuron expressing GCaMP6f (hereinafter, Mi1 > GC6f), plotted as the change in fluorescence relative to baseline (ΔF/F). **(F)** Linear filters represent each neuron’s dynamics by characterizing how they respond to preceding stimuli. Plotted filters correspond to Mi1 (Mi1 > GC6f, n = 68 flies), Tm3 (Tm3 > GC6f, n = 25 flies), Mi4 (Mi4 > GC6f, n = 15 flies), and CT1 (CT1 > GC6f, n = 17 flies). Linear filters are normalized to the maximum response of each fly’s mean filter. Lines are mean ± SEM. Neural response dynamics can be quantified by the filter’s half-rise (‘rise’), peak amplitude (‘peak’), and half-fall (‘fall’) times.

*Drosophila*’s motion detection circuits are anatomically and functionally well-characterized. In the fly eye, light intensity is first detected by photoreceptors before signals are split into ON and OFF pathways that detect light increments and decrements, respectively (Clark et al., 2011; Joesch et al., 2010; Silies et al., 2014). Within each pathway, interneurons delay and rectify visual signals (Arenz et al., 2017; Behnia et al., 2014; Strother et al., 2014; Yang et al., 2016) before synapsing onto the elementary direction-selective (DS) neurons of the ON and OFF pathways, T4 and T5 (Maisak et al., 2013). T4 and T5 neurons are classified into subtypes that respond preferentially to motion in one of four cardinal directions (Maisak et al., 2013). At least four types of interneurons, with different spatiotemporal response profiles, synapse onto T4 cells (Shinomiya et al., 2019; Takemura et al., 2017), which then integrate these signals to generate DS responses (Badwan et al., 2019; Gruntman et al., 2018; Haag et al., 2016; Leong et al., 2016; Salazar-Gatzimas et al., 2016; Strother et al., 2017). Output signals from T4 and T5 cells are then summed over space to guide visually-evoked behaviors (Creamer et al., 2018; Leonte et al., 2021; Maisak et al., 2013; Schilling and Borst, 2015).

Anatomical and physiological studies have suggested different models to explain how T4 cells detect the direction and speed of motion (Arenz et al., 2017; Badwan et al., 2019; Gruntman et al., 2018; Haag et al., 2016; Leong et al., 2016; Salazar-Gatzimas et al., 2018; Shinomiya et al., 2019; Strother et al., 2017; Zavatone-Veth et al., 2020), all of which depend on relative delays between signals at adjacent points in space (Figure 1A). In textbook versions of these models (Barlow and Levick, 1965; Hassenstein and Reichardt, 1956), the tuning of the motion detector to different velocities is fully determined by the relative delay in peak responses between two inputs (Figure 1A). Accordingly, changing the relative delay should predictably alter the tuning of DS signals. It is untested whether such delays are sufficient to explain how input neurons tune motion detection in *Drosophila*, or whether more complex temporal processing properties must be considered. More broadly, it remains unclear how DS circuits achieve selectivity for different speeds of motion. Neuro-modulators alter the tuning of motion detectors and the dynamics of their inputs (Arenz et al., 2017), but they act broadly and alter many properties, including the dynamics, of many neurons in the circuit (Strother et al., 2018). Thus, it also remains unknown how the dynamics of individual excitatory and inhibitory cell types contribute to downstream motion detection.

In this work, we altered the expression of specific membrane ion channels in four individual excitatory and inhibitory cell types in the fly motion detection circuit. We showed that these genetic manipulations of single cell types alter the dynamics of light responses in these neurons. Then, to test models of motion estimation, we asked how those manipulations of neural dynamics influence the tuning of downstream motion signals in T4 neurons. To do this, we manipulated ion channel expression in individual neuron types upstream of T4 neurons while measuring the responses of T4 to different speeds of visual motion. This resulted in altered tuning curves, showing how the different manipulations changed the sensitivity of T4 neurons to motion of different speeds. In the case of an interneuron that influences the ON and OFF motion pathways, we showed that these changes are also reflected in behavior. Last, we developed circuit models that are strongly constrained by anatomy and our measurements of response dynamics. We compared these models to our experimental data, and found that parallel, redundant excitatory and inhibitory inputs are required to explain our experimental data. Moreover, the full linear filtering properties of the inputs—rather than just delays—are necessary to reproduce our experimental observations. These results reveal how the timing of excitatory and inhibitory inputs generate motion signals and tune their sensitivity.

## Results

### Measuring the response dynamics of medulla neurons using stochastic visual stimuli

To investigate the role of individual interneurons in motion detection, we first measured the dynamic visual responses of inputs to T4 cells. We targeted four ON-cell types with anatomically identified synapses onto T4: Mi1, Tm3, Mi4, and CT1 (Figure 1B) (Takemura et al., 2017). Using *in vivo* two-photon microscopy, we recorded responses of these different cell types expressing the calcium indicator GCaMP6f (Chen et al., 2013) (Figure 1C), while their activity was driven by a stochastic, binary stimulus (Figure 1D). From these neural responses, we used standard methods (Chichilnisky, 2001) to extract the linear filters that best predicted the neuron’s response to the preceding stimulus (Figures 1E-F). While this method does not capture all the features of temporal processing, these filters can quantify many dynamical response properties of these neurons. For instance, a peak response that occurs after a short delay corresponds to a fast filter that represents a fast neural response to light signals. The filter shape also determines how much signal is passed at different temporal frequencies, with narrowly peaked filters transmitting more signal at high temporal frequencies. Consistent with previous findings, Mi1 dynamics were slower than Tm3 (Behnia et al., 2014), while both Mi1 and Tm3 dynamics were faster than Mi4 (Arenz et al., 2017; Strother et al., 2017) (Figure 1F). The dynamics of CT1 terminals were also consistent with previous measurements (Figure 1F) (Meier and Borst, 2019).

### Manipulating endogenous ion channel expression alters neural dynamics

After measuring the wildtype dynamics of Mi1, Tm3, Mi4, and CT1, we designed experiments to manipulate these cells by increasing or decreasing the expression of specific ion channels while co-expressing GCaMP6f to record the neuron’s response. We first tested how Mi1 dynamics were affected by knocking down several candidate ion channels, using either RNA interference (RNAi) or dominant-negative mutations (Figure S1). Based on these experiments, we chose to pursue manipulations using the channels *slowpoke* and *cacophony* because they had the largest effect sizes, are widely expressed in flies, and elicited opposing changes in Mi1 dynamics. We first manipulated the expression levels of *slo* (Elkins et al., 1986), a voltage-gated, Ca^2+^-activated K^+^ channel and ortholog of BK-type channels in vertebrates (Marty, 1981; Pallotta et al., 1981). *Slowpoke* is widely expressed in *Drosophila* neurons, including many visual neurons (Becker et al., 1995; Davis et al., 2018). It has an established role in modulating neural excitability and membrane conductance (Ford and Davis, 2014; Pattillo et al., 2001; Sun et al., 2004), and has relatively slow dynamics (Sah and Faber, 2002)—a property that makes it a candidate for helping induce the delays involved in *Drosophila* motion detection (Salazar-Gatzimas et al., 2016). The RNAi knock-down of *slo* (Perkins et al., 2015) slowed the dynamics of Mi1 slightly, demonstrating that *slowpoke* is necessary for wildtype dynamics (Figures 2A-B). If reduced *slo* slows the cell, we hypothesized that increased *slo* might speed it up. Indeed, when *slo* was over-expressed in Mi1, responses became faster, as quantified by faster filter peak and fall times (Figures 2C-D). Thus, manipulations of *slo* expression in Mi1 bi-directionally altered its dynamics.

**Figure 2.**
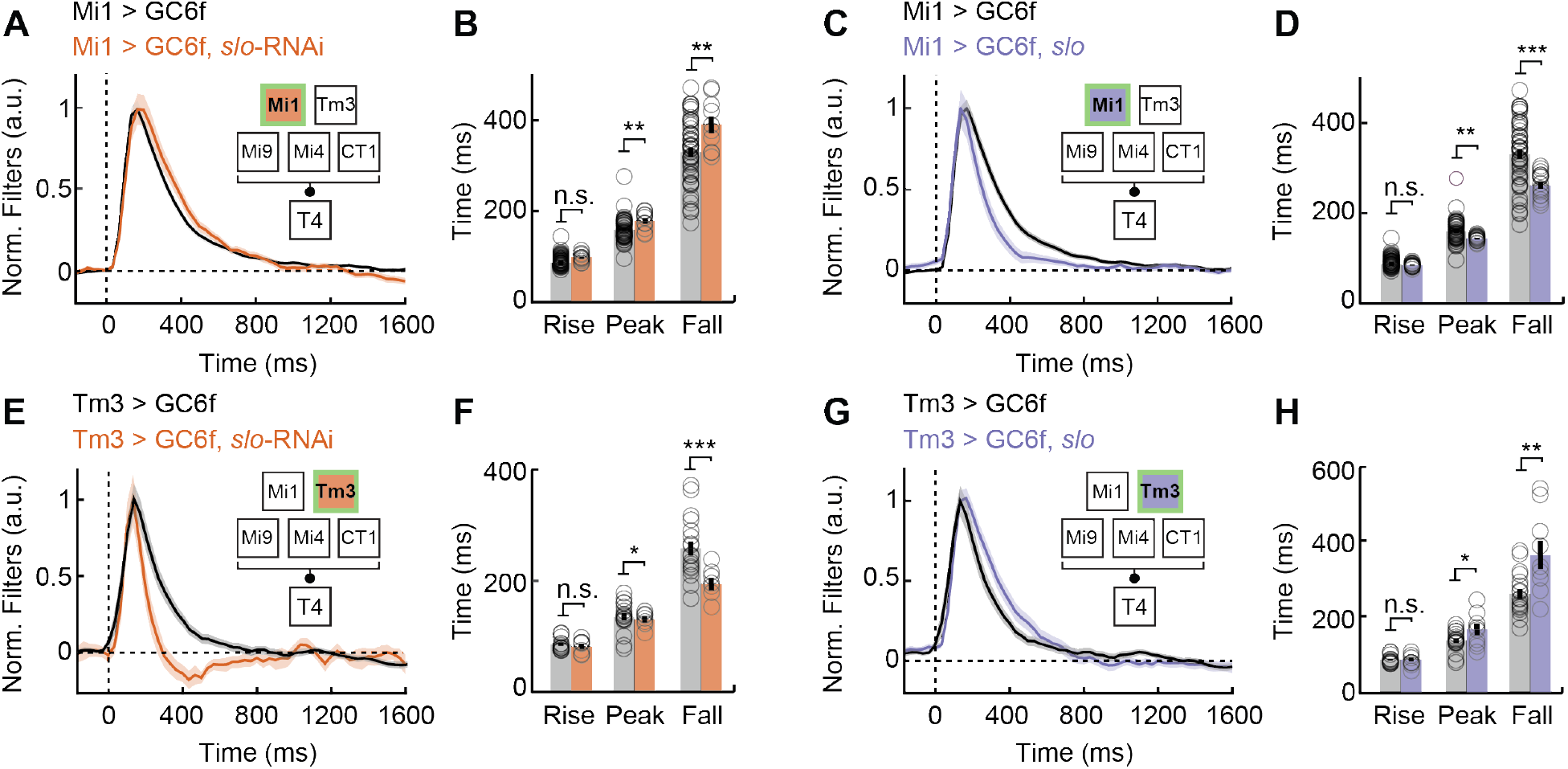
Cell-type-specific genetic manipulations of excitatory neurons Mi1 and Tm3 alter their dynamics. **(A)** Filters of Mi1 expressing *slo*-RNAi (Mi1 > GC6f, n = 68; Mi1 > GC6f, *slo*-RNAi, n = 19). Lines are mean ± SEM. **(B)** Half-rise (rise), peak, and half-fall (fall) times averaged across flies for the filters in (A). **(C-D)** As in (A-B), but with Mi1 over-expressing *slowpoke* (*slo*) (Mi1 > GC6f, *slo*, n = 16), compared to Mi1 native filter kinetics (Mi1 > GC6f, n = 68). **(E-F)** As in (A-B), but with Tm3 expressing *slo*-RNAi (Tm3 > GC6f, n = 25; Tm3 > GC6f, *slo*-RNAi, n = 19). **(G-H)** As in (A-B), but with Tm3 over-expressing *slo* (Tm3 > GC6f, n = 25; Tm3 > GC6f, *slo*, n = 8). **(I-J)** As in (A-B), but with Mi4 expressing an RNAi to knock-down *cacophony* (*cac*) (Mi4 > GC6f, n = 15; Mi4 > GC6f, *cac*-RNAi, n = 11). **(K-L)** As in (A-B), but with CT1 expressing *cac*-RNAi (CT1 > GC6f, n = 17; CT1 > GC6f, *cac*-RNAi, n = 11). (* p<0.05, ** p<0.01, *** p<0.001 by Wilcoxon signed-rank tests across flies.)

To investigate whether the role of *slo* generalized to other neurons, we performed the identical over-expression and RNAi knock-down experiments in Tm3 neurons (Figures 2E-H). Interestingly, each manipulation had the opposite effect in Tm3 as they had in Mi1. Expressing *slo*-RNAi in Tm3 resulted in faster responses, significantly reducing the filter fall time (Figures 2E-F), while over-expressing *slo* in Tm3 resulted in slower responses (Figures 2G-H). A second, distinct *slo*-RNAi construct (Dietzl et al., 2007) showed similarly strong effects on the response in Tm3, arguing against off-target effects for this large knock-down effect (Figure S2). The opposing results in our experiments are consistent with other distinct processing properties of Mi1 and Tm3, including their differing adaptation to stimulus contrast (Matulis et al., 2020) and their opposite responses to behavioral arousal (Strother et al., 2018). These experiments demonstrate that wildtype *slo* expression is required for both Mi1 and Tm3 wildtype dynamics, while the specific effect of manipulating *slo* expression appears to depend on the complement of channels expressed in the cell. Parallel experiments where the bacterial voltage-gated Na^+^ channel NaChBac (Nitabach et al., 2006) was expressed in either Mi1 or Tm3 cells also resulted in opposite changes in the dynamics of the two cell types (Figure S3). The changes in Mi1 and Tm3 dynamics were present in both dendrites and axon terminals, suggesting that they impact early stages of cellular processing (Figure S4).

To investigate whether these genetic manipulations affected membrane potential dynamics, we measured Mi1 and Tm3 voltage responses using Arclight (Jin et al., 2012) while using the manipulations that elicited the largest effects we observed with calcium indicators. Expressing *slo*-RNAi in Tm3 and NaChBac in Mi1 sped up each cell’s membrane potential response, consistent with our calcium measurements (Figure S5). This suite of manipulations in Mi1 and Tm3 cells did not strongly affect calcium response nonlinearities or filter amplitudes (Figure S6), suggesting that these manipulations do not strongly alter the basal physiological state of these neurons.

Next, we set out to manipulate the dynamics of the inhibitory neurons Mi4 and CT1. Mi4 has been anatomically (Takemura et al., 2013, 2017) and functionally (Strother et al., 2017) linked to T4, with other studies supporting its putative role as a delayed inhibitory input (Arenz et al., 2017; Gruntman et al., 2018). On the other hand, the role of CT1, an amacrine cell, in motion detection remains unknown, despite its shared characteristics with Mi4: it releases the inhibitory neurotransmitter GABA (Takemura et al., 2017), responds to local contrast increments (Meier and Borst, 2019), and synapses onto T4 with an anatomy that parallels Mi4 (Shinomiya et al., 2019). Due to the putative roles of Mi4 and CT1 as delayed inhibitory inputs, we sought to speed up their dynamics to determine how each cell type’s timing impacts T4 tuning.

Since *slo* over-expression and knock-down had opposite effects in Mi1 and Tm3, we used an alternative genetic manipulation that had the same effect on the dynamics of these two cell types. Knocking-down *cacophony* (*cac*), the voltage-gated Ca^2+^ α1 channel subunit, sped up both Mi1 and Tm3 filter dynamics (Figure S7). Similarly, when we used RNAi to knock-down *cac* in Mi4 and CT1, it made their responses significantly faster (Figures 3A-B). *Cac* knock-down in CT1 sped up its filter dynamics at terminals in both the medulla (Figures 3C-D) and the lobula (Figure S8). With this manipulation, the filter amplitudes of both Mi4 and CT1 responses were decreased, consistent with previous data using a gene excision method (Figure S6) (Fisher et al., 2017), but their nonlinearities showed relatively little change (Figure S6). These results demonstrate that *cac* expression is required to maintain Mi1, Tm3, Mi4, and CT1 wildtype calcium dynamics.

**Figure 3.**
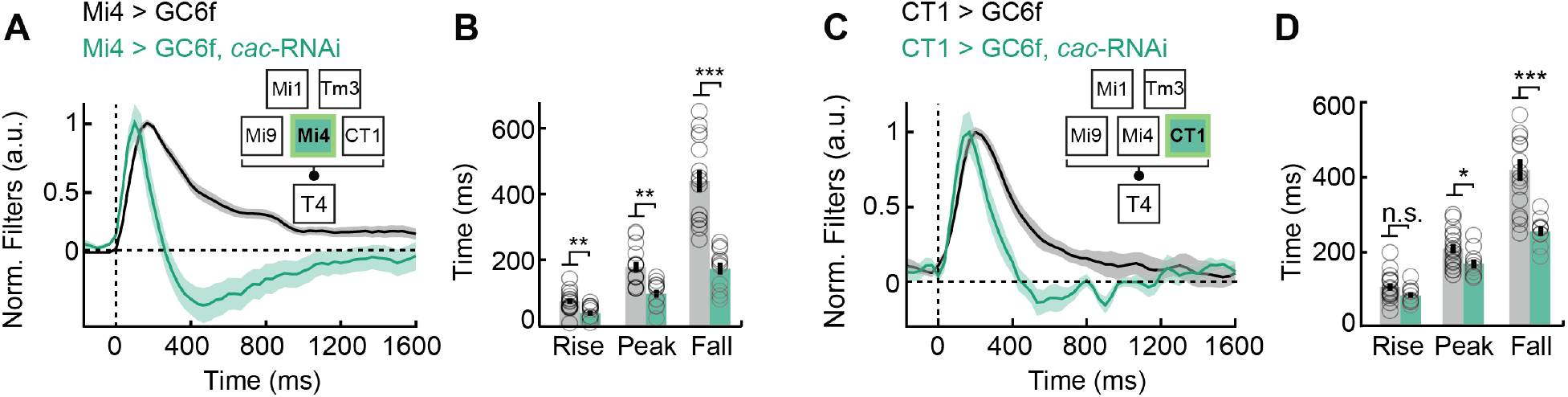
Cell-type-specific genetic manipulations of inhibitory neurons Mi4 and CT1 alter their dynamics. **(A)** Filters of Mi4 expressing an RNAi to knock-down *cacophony* (*cac*) (Mi4 > GC6f, n = 15; Mi4 > GC6f, *cac*-RNAi, n = 11). Lines are mean ± SEM. **(B)** Half-rise (rise), peak, and half-fall (fall) times averaged across flies for the filters in (A). **(C-D)** As in (A-B), but with CT1 expressing *cac*-RNAi (CT1 > GC6f, n = 17; CT1 > GC6f, *cac*-RNAi, n = 11). (* p<0.05, ** p<0.01, *** p<0.001 by Wilcoxon signed-rank tests across flies.)

### Excitatory and inhibitory input dynamics regulate T4 tuning to motion velocity

It is not surprising that manipulating membrane ion channel expression can alter response dynamics, but these manipulations enable us to interrogate how input dynamics drive downstream neural signals. We used these tools to investigate how T4 responses are determined by the dynamics of its excitatory and inhibitory inputs. To do this, we sped up or slowed down the dynamics of these inputs by expressing *slo*, *slo*-RNAi, or *cac*-RNAi, all while recording calcium responses in T4 cells (Figure 4–5). To measure the velocity tuning of T4, we presented periodic, white bars that rotated about the fly at different velocities (Figure 4A), and then compared T4 velocity sensitivity between manipulated conditions and controls. We recorded responses in T4 axons that responded to horizontal motion, and then combined responses across different preferred directions (PD) (Salazar-Gatzimas et al., 2016, 2018). As expected, T4 cells showed strong DS responses across the different velocities (Figure 4B). We plotted the tuning curve of each fly by averaging the responses over the 5 second presentation of each velocity (Figure 4C). These tuning curves peaked at around 32°/s. To summarize the speed tuning of these responses, we computed a response-weighted average that defines the curve’s center of mass on a log-velocity scale (Figure 4D, see Methods).

**Figure 4.**
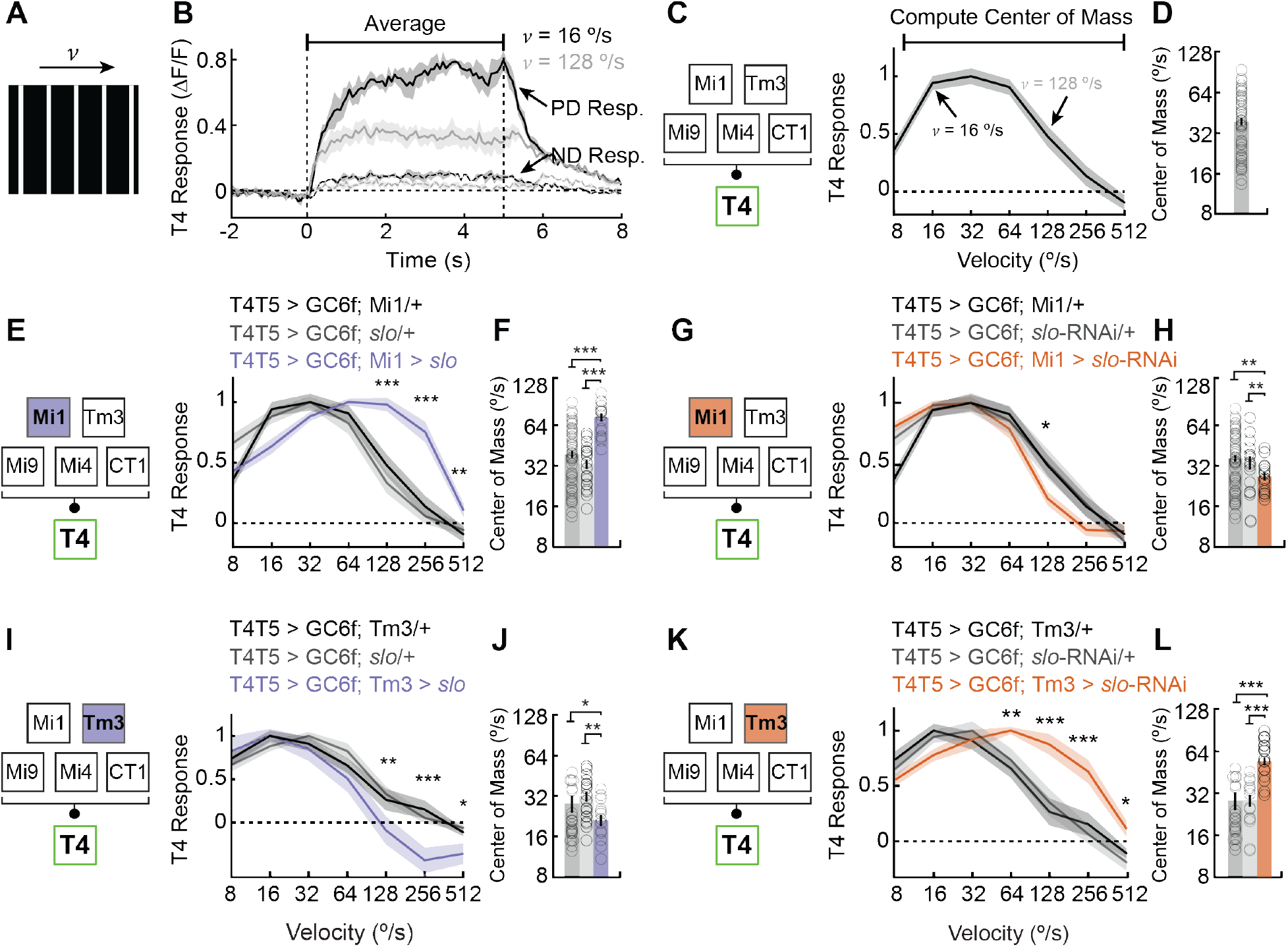
Genetic manipulations of excitatory inputs Mi1 and Tm3 dynamics alter T4 tuning. **(A)** The stimulus used to probe T4 tuning consists of white 5°-wide bars with 30° spacing rotating rightward and leftward at speeds between 8 and 512°/s. **(B)** Average T4 responses to white bars moving in the preferred direction (PD) and the null direction (ND) at 16°/s and 128°/s. Both PD and ND responses are averaged over the 5 second stimulus presentation window (n = 43 flies). **(C)** Example tuning curve computed from the raw response trace in (B). PD responses are shown, and each fly’s curve is normalized to its maximum response before averaging, depicted by black, horizontal bar. Curves shows mean and shading shows SEM. **(D)** The tuning curve’s log-velocity center of mass is a weighted average of the tuning curve shown in (C). Bars are mean ± SEM. **(E)** T4 tuning curves of flies over-expressing *slowpoke* (*slo*) in Mi1 (T4T5 > GC6f, Mi1 > *slo*, n = 9) compared to two genetic controls (T4T5 > GC6f; Mi1/+, n = 43 and T4T5 > GC6f; *slo*/+, n = 11). Lines are mean ± SEM. **(F)** Center of mass of T4 tuning curves from genotypes in (E). **(G-H)** As in (E-F), but for Mi1 expressing *slo*-RNAi (T4T5 > GC6f, Mi1 > *slo*-RNAi, n = 7) compared to two genetic controls (T4T5 > GC6f; Mi1/+, n = 43 and T4T5 > GC6f; *slo*-RNAi/+, n = 10). **(I-J)** As in (E-F), but for Tm3 over-expressing *slo* (T4T5 > GC6f, Tm3 > *slo*, n = 7) compared to two genetic controls (T4T5 > GC6f; Tm3/+, n = 11 and T4T5 > GC6f; *slo*/+, n = 11). **(K-L)** As in (E-F), but for Tm3 expressing *slo*-RNAi (T4T5 > GC6f, Tm3 > *slo*-RNAi, n = 12) compared to two genetic controls (T4T5 > GC6f; Tm3/+, n = 11 and T4T5 > GC6f; *slo*-RNAi/+, n = 10). (* p<0.05, ** p<0.01, *** p<0.001 by Wilcoxon signed-rank tests across flies. When there are two controls (in E, G, I, K), the reported significance is the larger of the comparisons to the two controls.)

**Figure 5.**
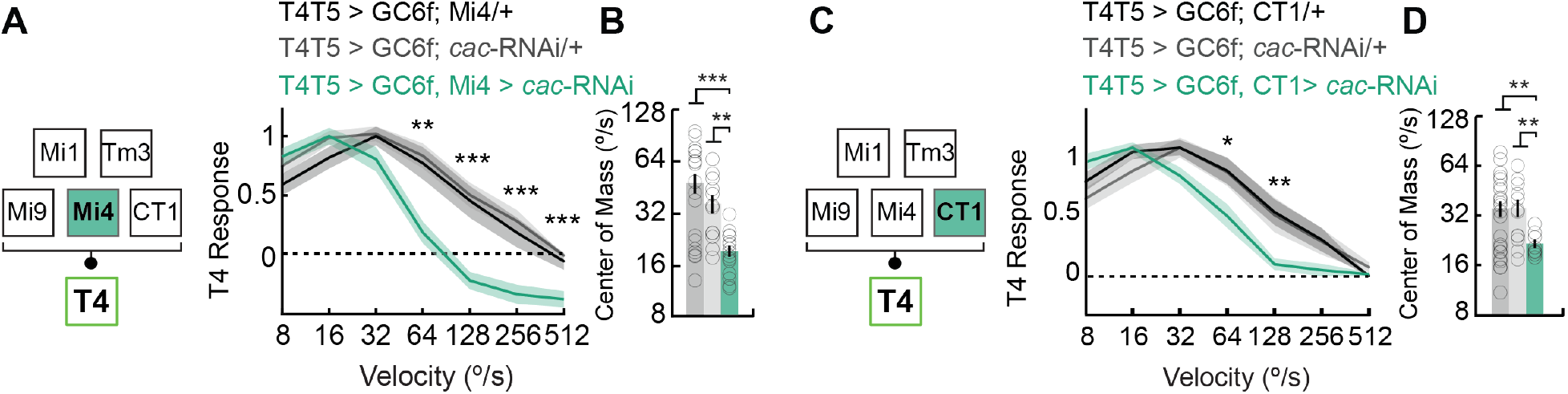
Genetic manipulations of inhibitory inputs Mi4 and CT1 dynamics alter T4 tuning. **(A)** T4 tuning curves of flies expressing an RNAi to knock-down *cacophony* (*cac*) in Mi4 (T4T5 > GC6f, Mi4 > *cac*-RNAi, n = 8) compared to two genetic controls (T4T5 > GC6f; Mi4/+, n = 12 and T4T5 > GC6f; *cac*-RNAi/+, n = 8). Lines are mean ± SEM. **(B)** Center of mass of T4 tuning curves from genotypes in (A). **(C-D)** As in (A-B), but for CT1 expressing *cac*-RNAi (T4T5 > GC6f, CT1 > *cac*-RNAi, n = 7) compared to two genetic controls (T4T5 > GC6f; CT1/+, n = 12 and T4T5 > GC6f; *cac*-RNAi/+, n = 8). (* p<0.05, ** p<0.01, *** p<0.001 by Wilcoxon signed-rank tests across flies. When there are two controls (in A, C), the reported significance is the larger of the comparisons to the two controls.)

We began by assessing the impact of Mi1 and Tm3 dynamics on T4 velocity tuning. If these two excitatory inputs serve as the non-delay inputs to T4 (Shinomiya et al., 2019), then speeding them up should lengthen relative delays in the circuit. The textbook model of circuit delays would predict that this should result in downstream motion signals that prefer slower stimuli (Figure 1A). To test this prediction, we sped up Mi1 dynamics by over-expressing *slo* (Figures 2A-B), and measured T4 responses. With this manipulation, we observed an increase in T4 sensitivity to bars moving at high speeds and a shift of the curve’s center of mass to higher velocities (Figures 4E-F). This change was opposite the prediction of the textbook model of circuit delays for motion detection. Conversely, slowing Mi1 by knocking-down *slo*, caused a small but significant decrease in sensitivity to high velocities (Figures 4G-H). The downstream consequences of manipulations to Tm3 dynamics paralleled those caused by altering Mi1 dynamics. When Tm3 was slowed down by *slo* over-expression, T4’s sensitivity to high velocities was reduced and the tuning curve’s center of mass shifted to slower velocities (Figures 4I-J). Likewise, when Tm3 dynamics were sped up by expressing *slo*-RNAi, T4 cells were significantly more sensitive to bars moving at high speeds (i.e., 64°/s-512°/s) (Figures 4K-L). In some cases, genetic manipulation of Mi1 and Tm3 altered T4 response amplitudes to the PD, but not to the null direction (ND) (Figure S9). In sum, speeding up or slowing down Mi1 or Tm3— two excitatory inputs—impacts T4 in a consistent fashion, but not as predicted by the textbook, delay-based model for motion detection.

We next assessed how altering the dynamics of the inhibitory inputs Mi4 and CT1—putative delay lines—affected T4 velocity tuning. Again, according to the textbook model for motion detection, making the delayed line faster should result in shorter relative delays, rendering the downstream motion detector more responsive to faster stimuli (Figure 1A). Therefore, we hypothesized that speeding up Mi4 or CT1 would result in T4 neurons that were more sensitive to faster velocities. Surprisingly, when we sped up Mi4 and CT1 by knocking down *cac*, we observed a significant increase in T4’s sensitivity to slower velocities (Figure 5A-D), contradicting the predictions of a simple, textbook model. CT1 has been anatomically implicated in T4 motion detection (Takemura et al., 2017) and it compartmentalizes signals that could potentially support local motion detection (Meier and Borst, 2019). However, there has been no functional evidence for its involvement. Our results show that the dynamics of CT1 are required for the tuning of T4. As with manipulations of Mi1 and Tm3, manipulating Mi4 and CT1 response dynamics did not substantially change T4 PD and ND response amplitudes (Figure S9). Interestingly, silencing Mi4 or CT1 with tetanus toxin did not result in changes in T4 tuning (Figure S10). This suggests that manipulating dynamics can reveal roles that are difficult to find using silencing experiments. In sum, these experiments show that speeding up Mi4 and CT1 responses significantly altered T4 velocity tuning in a similar fashion.

We wanted to test whether the tuning changes we observed in T4 were transmitted downstream to guide direction-selective behaviors in the fly. Therefore, we measured optomotor turning responses to the periodic, white bar stimulus we used to probe T4 tuning. We manipulated CT1 because it synapses onto both T4 and T5 neurons, which are both likely to be activated by our periodic stimulus. We hypothesized that expressing *cac*-RNAi in CT1 neurons would result in behavioral tuning changes matching the changes we observed in T4. Indeed, flies expressing *cac*-RNAi in CT1 were more sensitive to bars moving at lower velocities (Figure S11). These results reveal that (1) the tuning of T4 is transmitted to modulate fly turning behavior and (2) CT1 dynamics maintain native tuning to stimulus velocity in optomotor behavior.

### A data-driven model with parallel, delayed inhibitory inputs reproduces T4 velocity tuning

The simple, textbook model of circuit delays did not predict how altering the dynamics of excitatory (Mi1/Tm3) or inhibitory (Mi4/CT1) inputs changed T4 tuning. To better understand how the dynamical processing properties of these upstream neurons affects T4 responses, we compared our measurements to an anatomically-constrained synaptic model that incorporated the measured temporal filtering properties of the input neurons (Figure 6) (Badwan et al., 2019; Borst, 2018; Zavatone-Veth et al., 2020). This model consists of three, spatially-separated inputs that apply linear-nonlinear transformations to local visual signals (Zavatone-Veth et al., 2020). In this model, a central excitatory Mi1/Tm3-like ON input is flanked by an Mi9-like ND-offset OFF inhibitory input, and an Mi4-like PD-offset ON inhibitory input—all consistent with previous anatomical and functional data (Arenz et al., 2017; Gruntman et al., 2018; Strother et al., 2017; Takemura et al., 2017). We asked how this model responded to the periodic white bar stimulus used in T4 measurements (Figures 4–5 and 6A). To obtain data-driven filters for the inputs to this model, we first de-convolved calcium indicator dynamics from experimentally measured filters and then generated smooth filters by fitting with a parametric model (Figure 6B and S12, see Methods).

**Figure 6.**
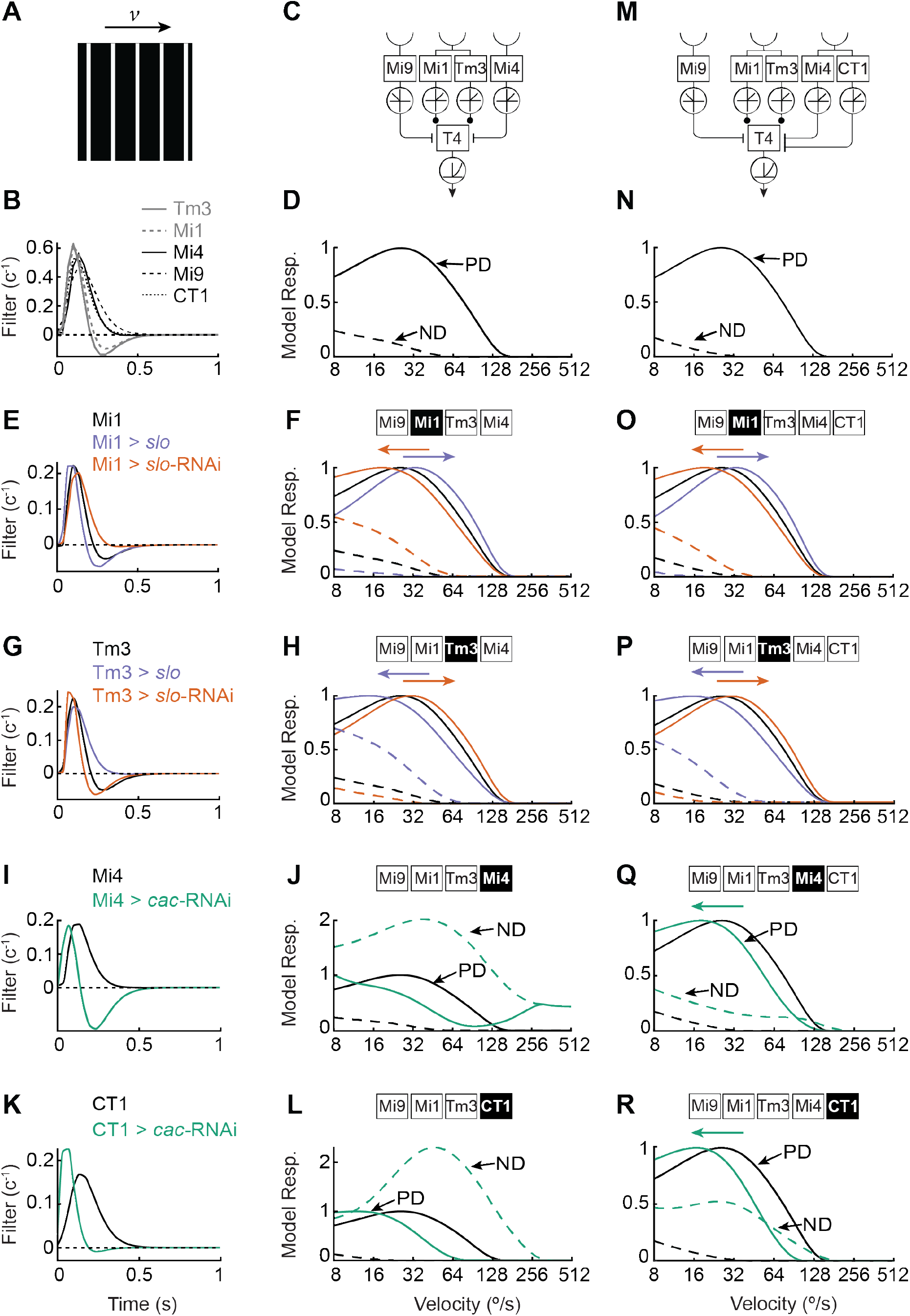
A synaptic model requires parallel, delayed inhibitory inputs to reproduce experimental results. **(A)** The experimental stimulus was used to simulate model responses. **(B)** Data-driven model filters were produced by de-convolving indicator dynamics from measured filters and then smoothing (see Methods). **(C)** Anatomically constrained synaptic model composed of three spatial inputs to T4: on the model’s null direction side, Mi9 is simulated as a delayed, OFF-responsive, inhibitory input; in the center, Mi1 and Tm3 share one spatial input and provide excitatory input; and on the model’s preferred direction side, Mi4 serves as a delayed, ON-responsive, inhibitory input. **(D)** The data-driven wildtype filters of each cell type were used to simulate the wildtype model’s response to the stimulus used in Figure 4–5 as it moved in preferred (PD) and null directions (ND) at different speeds. **(E-F)** As in (D), but with filters from wildtype Mi1, Mi1 over-expressing *slowpoke* (*slo*) (Mi1 > *slo*), and Mi1 expressing *slo*-RNAi (Mi1 > *slo*-RNAi). **(G-H)** As in (D), but with filters from wildtype Tm3, Tm3 over-expressing *slo* (Tm3 > *slo*), and Tm3 expressing *slo*-RNAi (Mi1 > *slo*-RNAi,). **(I-J)** As in (D), but with filters from wildtype Mi4 and Mi4 with *cacophony* (*cac*) knocked-down (Mi4 > *cac*-RNAi). **(K-L)** As in (D), but with filters from wildtype CT1 and CT1 expressing *cac*-RNAi (CT1 > *cac*-RNAi). **(M)** As in (C), but for an extended synaptic model with two parallel, delayed inhibitory inputs representing Mi4 and CT1. (**N**) The data-driven filters from (B) were used to simulate the model’s response in the presence of a parallel, delayed inhibitory input. **(O)** As in (N), but with the filters used in (E). **(P)** As in (N), but with the filters used in (G). **(Q)** As in (N), but with the filters used in (I). **(R)** As in (N), but with the filters used in (K).

To test how the excitatory Mi1 and Tm3 dynamics might alter tuning of T4 neurons in this model, we set up Mi1 and Tm3 as parallel linear-nonlinear synaptic inputs to T4 with a shared, central spatial receptive field (Figure 6C), again consistent with anatomical data (Takemura et al., 2017). Using these data-driven filters, we computed the model’s mean response to our periodic white bar stimulus rotating at different velocities (Figure 6D). The model’s PD response center of mass was ~32°/s, while its response to ND-moving bars was ~1/4 the amplitude of its PD response, both comparable to experimental measurements of T4 (Figure 4–5 and S9). Next, we simulated the model’s response when the Mi1 input used the data-driven filters for the experiments in which Mi1 expressed *slo* or *slo*-RNAi (Figure 6E). In the model, the faster dynamics of the Mi1 > *slo* filter shifted the model’s sensitivity toward faster velocities, while Mi1 > *slo*-RNAi filter shifted the sensitivity to slower velocities (Figure 6F), matching our experimental observations (Figure 4E-H). Similarly, the data-driven Tm3 > *slo* and Tm3 > *slo*-RNAi filters (Figure 6G) shifted the model’s sensitivity to slower and faster velocities, respectively (Figure 6H), also in agreement with our experiments (Figure 4I-L). These simulations make clear that the peak delay timing is not sufficient to qualitatively describe tuning changes; instead, the full bandpass properties of the filters are necessary to understand tuning of downstream motion detectors. When Mi1 and Tm3 become faster, they also pass more signal at high frequencies, resulting in the shift in tuning to higher velocities. This explanation is consistent with theoretical analyses of the simple Hassenstein-Reichardt correlator model (Egelhaaf and Borst, 1989; Reichardt, 1961), but these have never been directly tested. In all, these simulations show that the measured changes in the linear filtering properties of in Mi1 and Tm3 are sufficient to explain the consequent tuning changes measured in T4.

Next, we tested whether this model could explain our results when we manipulated the inhibitory Mi4 and CT1 input dynamics. When we substituted the Mi4 input with the Mi4 > *cac*-RNAi data-driven filter (Figure 6I), the model’s direction preference reversed, so that the response to periodic white bars moving in the former ND was *greater* than the response to those in the former PD (Figure 6J). This happened because the manipulated Mi4 delay line responds faster than the non-delay Mi1/Tm3 line. This simulation result is not supported by our experimental findings (Figure 4–5 and S13). We also asked whether the model could predict changes in T4 tuning if CT1, rather than Mi4, acted as the model’s delayed inhibitory input (Figures 6L-L). Exchanging the data-driven CT1 filter with that of CT1 > *cac*-RNAi (Figure 6K) also caused the model to reverse its direction preference (Figure 6L), a result similar to the Mi4 result and inconsistent with our T4 measurements of this manipulation (Figure 5).

These two failures of the initial model caused us to revise it. We created a new model in which Mi4 and CT1 both act as parallel, delayed, inhibitory inputs sharing the same spatial receptive field (Figure 6M), a proposal consistent with anatomy (Takemura et al., 2017). Using data-driven filters, this model architecture produced a velocity tuning curve that qualitatively resembled that of the previous model (Figure 6N). Similarly, adding the parallel, delayed inhibitory input did not change the model’s response to perturbations of the Mi1 or Tm3 inputs (using the data-driven filters corresponding to wildtype, *slo* over-expression, and *slo*-RNAi expression) (Figure 6O-P). However, in this model, when we exchanged the Mi4 or CT1 wildtype filters with the data-driven filters for Mi4 > *cac*-RNAi (Figure 6I) or CT1 > *cac*-RNAi (Figure 6K), the model’s direction preference remained intact (Figure 6Q-R). In the case of Mi4 > *cac*-RNAi, the model’s sensitivity shifted towards slower moving bars (Figure 6Q). In the case of CT1 > *cac*-RNAi, there was a similar shift in the sensitivity towards lower velocities (Figure 6R). Both cases matched the changes observed in T4 tuning (Figures 5A-B and 5C-D). Therefore, this revised synaptic model is sufficient to account for the changes in tuning of T4 when inhibitory Mi4 and CT1 dynamics are altered.

These results did not depend strongly on the details of the model. For instance, tuning shifts remained consistent when we replaced the data-driven filters with synthetic high- and low-pass filters (Figure S14). Our manipulations of Mi4 and CT1 both sped up the filters and also reduced their amplitudes (Figures 3A-D and S6), but simulations including both effects roughly matched those in which we altered only the filtering dynamics (Figure S15). In contrast, including only the reduction in amplitude in Mi4 or CT1, without the change in dynamics, resulted in tuning changes in T4 that were in the opposite direction of what we observed experimentally (Figure S15).

## Discussion

Overall, this research provides causal evidence for how the dynamics of four known input interneurons to T4—Mi1, Tm3, Mi4, and CT1—influence motion computation. First, we showed that ion channel expression levels regulate neural response dynamics. Specifically, we identified two membrane ion channels whose expression is required for the wildtype dynamics of various cell types. Next, we showed that manipulating the dynamics of single inputs alters T4 velocity tuning. The response dynamics of excitatory and inhibitory neuron types are combined to jointly tune T4 sensitivity to different velocities. These experimental observations of T4 tuning under different input manipulations are not explained by textbook models of motion detection that consider only the delays of inputs. Instead, the full, filtering properties of the filters are necessary to predict our experimental results. Finally, we showed that a data-constrained synaptic model for T4 reproduces our findings only when two delayed inhibitory inputs from Mi4 and CT1 are in parallel.

### Neurons can control their response dynamics by regulating ion channel expression

Studies have suggested many properties by which networks of neurons may regulate their processing dynamics, from conduction delays (Egger et al., 2020) and synaptic dynamics (Alabi and Tsien, 2012) to feedback and lateral circuit interactions (Drinnenberg et al., 2018). Our findings highlight how active membrane channel expression controls cellular response dynamics, and in turn regulate how circuit computations are tuned. In particular, we identified two ion channels—*slowpoke* and *cacophony*—that are critical to maintaining the native response dynamics of four input interneurons in the fly’s motion detection circuit (Figure 2–3 and S7). The four input interneurons we studied—Mi1, Tm3, Mi4, and CT1—use membrane channel expression to impose additional delays in signals and control their dynamics (Figure 2–3). It is not surprising that manipulating ion channel expression affects neural dynamics. In fact, ion channel expression has been shown to regulate neural dynamics in other *Drosophila* studies (Groschner et al., 2018; Gür et al., 2020), as well as with some timing mechanisms in vertebrate motor control circuits, which can rely on axonal conductance properties to coordinate activity (Egger et al., 2020). However, the method we used in this study to manipulate the dynamics of *individual* neuron types is a powerful tool for manipulating and dissecting circuit function. For neurons and circuits, regulating ion channel expression provides a flexible way to control their dynamics and circuit computations.

Neurons and circuits have homeostatic mechanisms that regulate membrane channel expression to ensure stable network function (Marder and Goaillard, 2006). Moreover, there are likely many channel expression patterns in a cell that could achieve similar response dynamics (Prinz et al., 2004). The interneurons we manipulated here are potentially under homeostatic control (Davis, 2006), yet our experiments successfully manipulated their dynamics. This suggests that homeostatic regulation is imperfect in these cells, as it relates to response dynamics, or that the dynamics are not being actively controlled by homeostatic mechanisms. The possibility of homeostatic regulation also warrants some caution in interpreting results: the misexpression of certain genes creates phenotypes in response dynamics, but those gene products are not necessarily the channels responsible for altering neural dynamics, since many channels could change in abundance or function. The opposite, bidirectional effects of manipulating *slo* in Mi1 and Tm3 also make it probable that dynamics are controlled by a complex interplay of channels that are different between these two neurons. The differences observed in Mi1 and Tm3 responses to *slo* manipulations are also consistent with experimental findings in vertebrates, where manipulating a potassium channel may either increase or decrease excitability, depending on the neuron type (Quraishi et al., 2019; Yang et al., 2007).

Manipulating cellular expression patterns to alter neural dynamics offers a circuit dissection tool that complements genetically encoded silencing methods, which have served as a primary tool for understanding circuit function (Luo et al., 2018). Interestingly, these manipulations revealed roles for Mi4 and CT1 in tuning motion detection that silencing did not (Figure 5 and S10). By altering neural properties but not silencing the neurons, these experiments act somewhat like activation experiments. That is, they alter neural activity as a function of on-going responses, and show that this changed activity is sufficient to affect different properties of the circuit.

### Excitatory and inhibitory input dynamics jointly control velocity sensitivity

In this research, we developed a protocol that allowed us to genetically manipulate individual inputs to T4 while simultaneously measuring the impact on T4 velocity tuning. Using this protocol, we demonstrated how perturbing the dynamics of Mi1, Tm3, Mi4, and CT1 each changed T4 sensitivity to stimulus velocity (Figure 4–5). Thus, this work reveals that each of these neuron types *individually* contributes to tuning velocity sensitivity in T4, while the dynamics of both excitatory and inhibitory inputs *jointly* control the tuning of T4. Moreover, although prior work has suggested that the amacrine cell CT1 could be involved in T4 function (Meier and Borst, 2019; Shinomiya et al., 2019; Takemura et al., 2017), our results demonstrate that its responses tune T4 motion detection. Last, T4 and T5 are required for rotational optomotor behaviors (Maisak et al., 2013), but it remained unknown how their tuning contributed to behavioral responses. Our manipulations showed that behavioral tuning changed in the same direction as T4 tuning (Figure S11).

The control of motion detector tuning by both excitatory and inhibitory dynamics may extend to motion detection circuits in mouse and other vertebrates. For instance, in mouse, both starburst and amacrine cells, as well as cortical DS cells, receive excitatory inputs with differential delays (Baden et al., 2013; Kim et al., 2014; Lien and Scanziani, 2018). These delays appear critical to direction-selectivity and could be, in part, generated by differential expression of active ion channels. Moreover, starburst and cortical cells receive direct and indirect inhibition from neighboring cells, and our results suggest that the dynamics of this inhibition could tune the velocity sensitivity of these cells. Last, DS retinal ganglion cells receive excitatory inputs from bipolar cells and directional inhibition from starburst cells (Demb and Singer, 2015). Our results suggest that the dynamics of *both* the excitation and the inhibition control the sensitivity of these cells to velocity.

### Manipulating single-neuron-type response dynamics to constrain circuit models

Our genetic manipulations of Mi1, Tm3, Mi4, and CT1 while recording T4 provide sensitive tests of models for motion detection in *Drosophila*. Our experimental and theoretical results (Figure 3–6) suggest that the excitatory and inhibitory interneurons tested play redundant roles in T4 tuning. This redundancy is consistent with neural anatomy, in which these two pairs of neurons receive input from similar points in space (Takemura et al., 2017). The redundancy is also consistent with the result that T4 largely maintains direction-selectivity even when its inputs are individually silenced (Strother et al., 2017). In addition, our data establish that the tuning of local motion detectors cannot be predicted by examining relative delays alone. Rather, our model suggests that it depends on detailed linear filtering properties of input neurons (Figure 4–5)—a hitherto untested theoretical result (Egelhaaf and Borst, 1989; Reichardt, 1961). The circuit simulations suggest that, although this circuit has many feedback and lateral connections (Takemura et al., 2013, 2017), a feedforward synaptic model can reproduce the tuning properties resulting from our manipulations of input dynamics.

Previous work has shown that modulating channel expression can determine network dynamics (Schulz et al., 2006), but this work shows how changes in channel expression in single neuron types can influence neural computation. More generally, because neural circuits ubiquitously integrate excitatory and inhibitory inputs, our results show how the dynamical responses of neural inputs are critical to understanding circuit computations. Beyond vision and motion detection, dynamics are central to many neural computations. For example, in auditory systems, interaural timing is crucial to localizing sounds (Grothe et al., 2010; Jeffress, 1948; Knudsen and Konishi, 1978, 1979), while in olfactory systems, the dynamics of odor responses facilitates odor discrimination (Laurent, 2002; Mazor and Laurent, 2005). Learning and synaptic plasticity also rely on the relative timing of neural activity (Dan and Poo, 2004), and motor control depends on the precise relative timing of neural signals (Churchland et al., 2012; Long et al., 2010). It will be interesting to investigate how the response timing of individual neurons in these systems drives circuit responses, and how response timing itself is influenced by the complement of membrane ion channels. Our results emphasize how expression of active channels in single excitatory and inhibitory neuron types can tailor neural computations in broader circuits.

## Materials and Methods

### Fly Strains and Husbandry

Non-virgin female flies, grown on dextrose-based food, were used for all experiments. All flies were staged on CO_2_ 12-24 hours after eclosion, and recordings were performed between 24 and 48 hours after staging. Both experimental and control flies used for imaging experiments were grown in incubators set to 25°C. All genotypes used are listed in **Table S1**, parental strains are listed in **Table S2**.

**Table S1.** Experimental Genotypes

| Abbreviation | Genotype |
| --- | --- |
| Mi1 > GC6f | +; UAS-GCaMP6f/+; R19F01-Gal4/+ |
| Mi1 > GC6f, *slo* | w, UAS-*slo*/+; UAS-GCaMP6f /+; R19F01-Gal4/+ |
| Mi1 > GC6f, NaChBac | w/+; UAS-GCaMP6f /+; R19F01-Gal4/UAS-NaChBac |
| Mi1 > GC6f, *slo*-RNAi | +; UAS-GCaMP6f/UAS-*slo*-RNAi; R19F01-Gal4/+ |
| Mi1 > GC6f, *GluClal*-RNAi | +; UAS-GCaMP6f/UAS-*GluClal*-RNAi; R19F01-Gal4/UAS-Dcr-2 |
| Mi1 > GC6f, *nAchRa1*-RNAi | w/+; UAS-GCaMP6f/UAS-*nAchRa1*-RNAi; R19F01-Gal4/+ |
| Mi1 > GC6f, *para*-RNAi | w/+; UAS-GCaMP6f/+; R19F01-Gal4/*para*-RNAi |
| Mi1 > GC6f, *Sh*-DN | w/+; UAS-GCaMP6f/UAS-*Sh*-DN; R19F01-Gal4/+ |
| Mi1 > GC6f, *eag*-DN | w/+; UAS-GCaMP6f/UAS-*eag*-DN; R19F01-Gal4/+ |
| Tm3 > GC6f | +; UAS-GCaMP6f /+; R13E12-Gal4/+ |
| Tm3 > GC6f, *slo* | w, UAS-*slo*/+; UAS-GCaMP6f /+; R23E12-Gal4/+ |
| Tm3 > GC6f, NaChBac | w/+; UAS-GCaMP6f /+; R13E12-Gal4/UAS-NaChBac |
| Tm3 > GC6f, *slo*-RNAi | +; UAS-GCaMP6f/UAS-*slo*-RNAi; R13E12-Gal4/+ |
| Mi4 > GC6f | w/+; UAS-GCaMP6f /R48A07-AD; R79H02-DBD/+ |
| Mi4 > GC6f, *cac*-RNAi | w/+; UAS-GCaMP6f/R48A07-AD; R79H02-DBD/UAS-*cac*-RNAi |
| CT1 > GC6f | w/+; UAS-GCaMP6f/R65E11-AD; R20C09-DBD/+ |
| CT1 > GC6f, *cac*-RNAi | w/+; UAS-GCaMP6f/R65E11-AD; R20C09-DBD/UAS-*cac*-RNAi |
| Tm3 > GC6f, *slo*-RNAi (v2) | +; UAS-GCaMP6f /UAS-*slo*-RNAi; R13E12-Gal4/+ (VDRC) |
| Mi1 > ArcLD | +; UAS-ArcLight/+; R19F01-Gal4/+ |
| Mi1 > ArcLD, NaChBac | w/+; UAS-ArcLight/+; R19F01-Gal4/UAS-NaChBac |
| Tm3 > ArcLD | +; UAS-ArcLight/+; R13E12-Gal4/+ |
| Tm3 > ArcLD, *slo*-RNAi | +; UAS-ArcLight/UAS-*slo*-RNAi; R19F01-Gal4/+ |
| Mi1 > GC6f, *cac*-RNAi | w/+; UAS-GCaMP6f /+; R19F01-Gal4/UAS-*cac*-RNAi |
| Tm3 > GC6f, *cac*-RNAi | w/+; UAS-GCaMP6f /+; R13E12-Gal4/UAS-*cac*-RNAi |
| T4T5 > GC6f, Mi1/+ | +; LexAOp-GCaMP6f, R42F06-LexA/+; R19F01-Gal4/+ |
| T4T5 > GC6f, *slo*/+ | w, UAS-*slo*/+; LexAOp-GCaMP6f, R42F06-LexA/+; + |
| T4T5 > GC6f, Mi1 > *slo* | w, UAS-*slo*/+; LexAOp-GCaMP6f, R42F06-LexA/+; R19F01-Gal4/+ |
| T4T5 > GC6f, *slo*-RNAi/+ | +; LexAOp-GCaMP6f, R42F06-LexA/UAS-*slo*-RNAi; + |
| T4T5 > GC6f, Mi1 > *slo*-RNAi | +; LexAOp-GCaMP6f, R42F06-LexA/UAS-*slo*-RNAi; R19F01-Gal4/+ |
| T4T5 > GC6f, Tm3/+ | +; LexAOp-GCaMP6f, R42F06-LexA/+; R13E12-Gal4/+ |
| T4T5 > GC6f, Tm3 > *slo* | w, UAS-*slo*/+; LexAOp-GCaMP6f, R42F06-LexA/+; R13E12-Gal4/+ |
| T4T5 > GC6f, Tm3 > *slo*-RNAi | +; LexAOp-GCaMP6f, R42F06-LexA/UAS-*slo*-RNAi; R13E12-Gal4/+ |
| T4T5 > GC6f, Mi4/+ | w/+; LexAOp-GCaMP6f, R42F06-LexA/R48A07-AD; R79H02-DBD /+ |
| T4T5 > GC6f, *cac*-RNAi/+ | w/+; LexAOp-GCaMP6f, R42F06-LexA/ +; UAS-*cac*-RNAi/+ |
| T4T5 > GC6f, Mi4 > *cac*-RNAi | w/+; LexAOp-GCaMP6f, R42F06-LexA/ R48A07-AD; R79H02-DBD /UAS-*cac*-RNAi |
| T4T5 > GC6f, CT1/+ | w/+; LexAOp-GCaMP6f, R42F06-LexA/R65E11-AD; R20C09-DBD/+ |
| T4T5 > GC6f, CT1 > *cac*-RNAi | w/+; LexAOp-GCaMP6f, R42F06-LexA/R65E11-AD; R20C09-DBD /UAS-*cac*-RNAi |
| T4T5 > GC6f, TNT/+ | w/+; LexAOp-GCaMP6f, R42F06-LexA/UAS-TNT; + |
| T4T5 > GC6f, Mi4 > TNT | w/+; LexAOp-GCaMP6f, R42F06-LexA/ R48A07-AD; R79H02-DBD /UAS-TNT |
| T4T5 > GC6f, CT1 > TNT | w/+; LexAOp-GCaMP6f, R42F06-LexA/R65E11-AD; R20C09-DBD /UAS-TNT |
| Mi9 > GC6f | w/+; UAS- GCaMP6f /R48A07-AD; VT046779-DBD/+ |
| T4T5/+ | +; R42F06-Gal4; + |
| TNT/+ | w/+; UAS-TNT/+; + |
| T4T5 > TNT | w/+; R42F06-Gal4/UAS-TNT; + |

**Table S2.** Parental Strains

| Genotype | Identifier | Source |
| --- | --- | --- |
| *D. melanogaster* Mi1: +; +; R19F01-Gal4 | 48852 | [(Strother et al., 2014)]; BDSC |
| *D. melanogaster* Mi4: +; R48A07-AD; R79H02-DBD | SS00316 | [(Strother et al., 2017)]; BDSC |
| *D. melanogaster* Tm3: +; +; R13E12-Gal4 | 48569 | [(Behnia et al., 2014)]; BDSC |
| *D. melanogaster* GCaMP6f: +; UAS-GC6f; + | 42747 | [(Chen et al., 2013)]; BDSC |
| *D. melanogaster* T4T5: +; R42F06-LexA; + | 54203 | BDSC |
| *D. melanogaster* GCaMP6f: +; LexAOp-GC6f; + | 44277 | BDSC |
| *D. melanogaster* TNT: +; UAS-TNT; + | 28838 | [(Sweeney et al., 1995)]; BDSC |
| *D. melanogaster slo*-RNAi: +; UAS-*slo*-RNAi (TRiP) | 55405 | [(Perkins et al., 2015)]; BDSC |
| *D. melanogaster slo*-RNAi v2: +; UAS-*slo*-RNAi | 104421 | [(Dietzl et al., 2007)]; VDRC |
| *D. melanogaster slo*: w, UAS-*slo*; +; + | Gift from Dr. N. S. Atkinson, The University of Texas at Austin, Austin, TX, USA | [(Nitabach et al., 2006)] |
| *D. melanogaster* NaChBac: w; UAS-NaChBac | 9468 | [(Nitabach et al., 2006; Shafer and Yao, 2014)]; BDSC |
| *D. melanogaster cac*-RNAi: yv; UAS-*cac*-RNAi | 27244 | [(Perkins et al., 2015)]; BDSC |
| *D. melanogaster GluClal*-RNAi: +; UAS-*GluClal*-RNAi | 105754 | [(Dietzl et al., 2007)]; VDRC |
| *D. melanogaster* UAS-Dcr-2: w; +; UAS-Dcr-2 | 24651 | [(Hartwig et al., 2008)]; BDSC |
| *D. melanogaster nAchRal*-RNAi: +; +; UAS-*nAchRal*-RNAi | 28688 | [(Perkins et al., 2015)]; BDSC |
| *D. melanogaster para*-RNAi: +; +; UAS-*para*-RNAi | 33923 | [(Perkins et al., 2015)]; BDSC |
| *D. melanogaster Sh*-DN: w; UAS-*Sh*-DN | Gift from Dr. Haig Keshishian, Yale University, New Haven, CT, USA | [(Gisselmann et al., 1989)] |
| *D. melanogaster eag*-DN: w; UAS-*eag*-DN | 8187 | [(Hartwig et al., 2008)]; BDSC |
| *D. melanogaster* ArcLight: w; UAS-ArcLight/CyO | 51057 | BDSC |
| *D. melanogaster* CT1: w; R65E11-AD; R20C09-DBD | SS01001 | [(Takemura et al., 2017)] |
| *D. melanogaster* Mi9: w; R48A07-AD; VT046779-DBD | SS00316 | [(Strother et al., 2017)] |

### Visual stimuli

Stimuli for imaging experiments were generated using custom code written in Matlab (The MathWorks, Natick, MA) and PsychToolBox (Brainard, 1997; Kleiner et al., 2007; Pelli, 1997). Both were projected with digital light projectors (Texas Instruments) onto panoramic screens surrounding the fly as described previously (Creamer et al., 2019). Stimulus frames were presented at an update rate of 180 Hz, and stimuli were presented in green light with a mean intensity of ~100 cd/m^2^. To minimize stimulus bleed-through onto microscope photomultiplier tubes (PMTs), the projector light was filtered with two 565/24 (centers/FWHM) filters in series (Semrock, Rochester, NY, USA). All visual stimuli presented in the experiments are listed in **Table S3**.

**Table S3.**
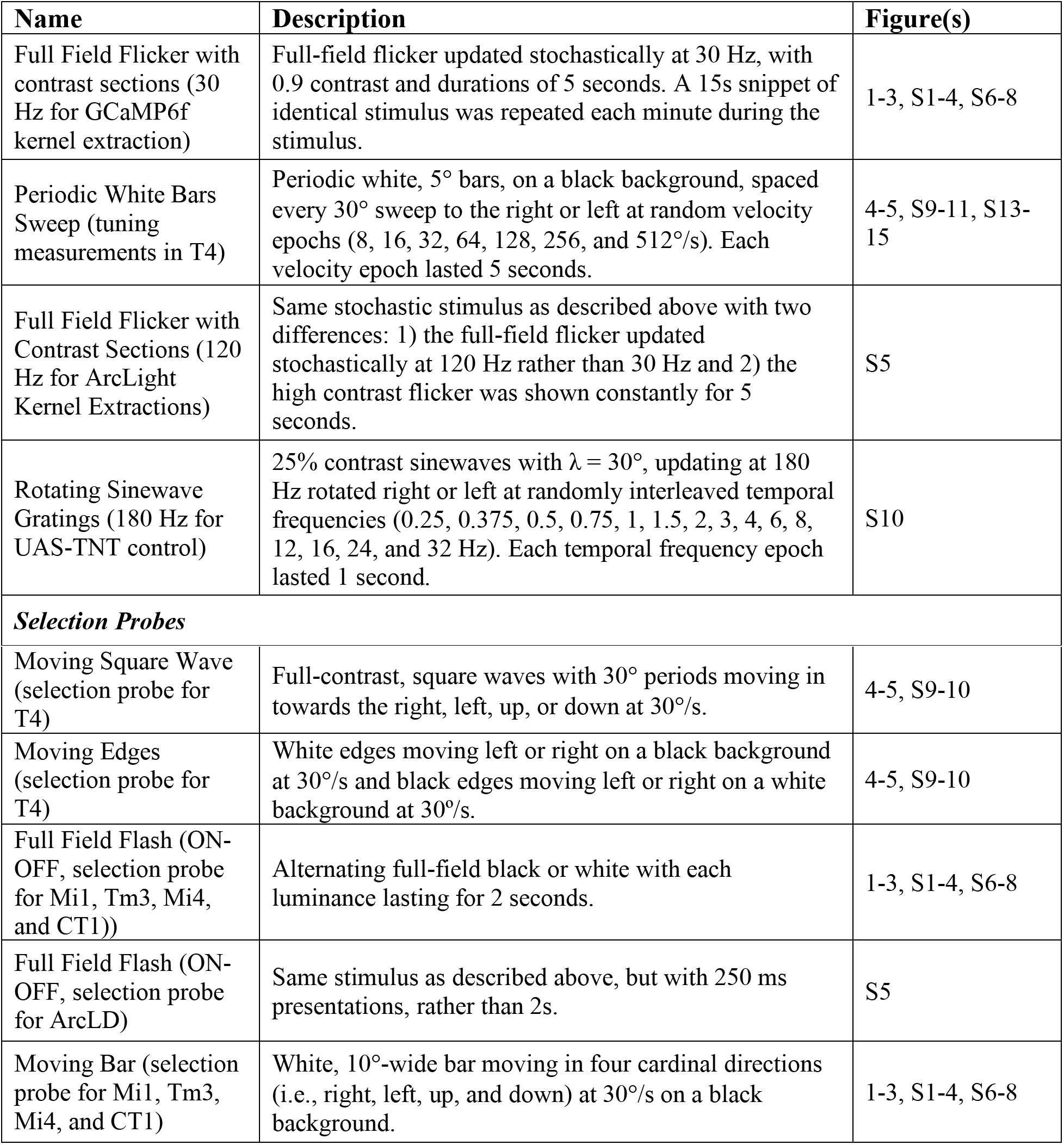
Visual Stimuli

| Name | Description | Figure(s) |
| --- | --- | --- |
| Full Field Flicker with contrast sections (30 Hz for GCaMP6f kernel extraction) | Full-field flicker updated stochastically at 30 Hz, with 0.9 contrast and durations of 5 seconds. A 15s snippet of identical stimulus was repeated each minute during the stimulus. | 1-3, S1-4, S6-8 |
| Periodic White Bars Sweep (tuning measurements in T4) | Periodic white, 5° bars, on a black background, spaced every 30° sweep to the right or left at random velocity epochs (8, 16, 32, 64, 128, 256, and 512°/s). Each velocity epoch lasted 5 seconds. | 4-5, S9-11, S13-15 |
| Full Field Flicker with Contrast Sections (120 Hz for ArcLight Kernel Extractions) | Same stochastic stimulus as described above with two differences: 1) the full-field flicker updated stochastically at 120 Hz rather than 30 Hz and 2) the high contrast flicker was shown constantly for 5 seconds. | S5 |
| Rotating Sinewave Gratings (180 Hz for UAS-TNT control) | 25% contrast sinewaves with $\lambda = 30^\circ$, updating at 180 Hz rotated right or left at randomly interleaved temporal frequencies (0.25, 0.375, 0.5, 0.75, 1, 1.5, 2, 3, 4, 6, 8, 12, 16, 24, and 32 Hz). Each temporal frequency epoch lasted 1 second. | S10 |
| **Selection Probes** |  |  |
| Moving Square Wave (selection probe for T4) | Full-contrast, square waves with 30° periods moving in towards the right, left, up, or down at 30°/s. | 4-5, S9-10 |
| Moving Edges (selection probe for T4) | White edges moving left or right on a black background at 30°/s and black edges moving left or right on a white background at 30°/s. | 4-5, S9-10 |
| Full Field Flash (ON-OFF, selection probe for Mi1, Tm3, Mi4, and CT1)) | Alternating full-field black or white with each luminance lasting for 2 seconds. | 1-3, S1-4, S6-8 |
| Full Field Flash (ON-OFF, selection probe for ArcLD) | Same stimulus as described above, but with 250 ms presentations, rather than 2s. | S5 |
| Moving Bar (selection probe for Mi1, Tm3, Mi4, and CT1) | White, 10°-wide bar moving in four cardinal directions (i.e., right, left, up, and down) at 30°/s on a black background. | 1-3, S1-4, S6-8 |

### Two-Photon Imaging Protocol

Fluorescent activity of labelled neurons was recorded using two-photon scanning fluorescence microscopy. Flies were anesthetized on ice and mounted onto stainless steel shim holders. Using UV-cured epoxy, we fixed the anterior rim of their heads to the holder. We surgically removed the posterior cuticle and trachea of the right or left eye. The flies’ brains were covered by oxygenated sugar-saline solution (Wilson et al., 2004). The metal holder was placed above a box of panoramic screens (Creamer et al., 2019) under a Scientifica two-photon microscope. The panoramic screens onto which the visual stimuli were projected subtended 270° in azimuth and 69° in elevation. Fluorophores were excited with a Spectra-Physics MaiTai eHP laser set to 930 nm wavelength and with power at the sample less than or equal to ~30 mW. Using ScanImage (Pologruto et al., 2003), images were acquired at approximately 13 Hz. To prevent undesirable bleed-through from the visual stimulus, the input to the PMT was filtered with a 512/25 and a 514/30 (center/FWHM) filter in series (Semrock, Rochester, NY, USA). All data were processed and analyzed using custom MATLAB code.

### Imaging Data Analysis: ROI Identification

For Mi1, Tm3, Mi4, and CT1 recordings, regions of interest (ROIs) were identified by hand to encompass one neuron per ROI. For Mi1 recordings, layers M1, M5, and M9/10 were analyzed, while for Tm3, layers M1, M4/5, and M9/10 were analyzed. For Mi4 recordings, layers M2/3/4 and M8/9 were analyzed. For CT1, terminals were recorded in the medulla M10 layer and in lobula layer L1. T4 axon terminals were recorded in the lobula plate, where we ran a watershed algorithm over the mean acquisition image to extract ROIs based on the baseline fluorescent intensity. For all, when low signal-to-noise ratio impeded identifying ROIs, we computed correlations in intensity over the movie between each pixel and its neighboring pixels, and used that ‘correlation image’ to define the boundaries of ROIs.

### Imaging Data Analysis: ROI Analysis

For each ROI, ΔF/F was computed with methods previously described (Salazar-Gatzimas et al., 2016). The baseline fluorescence *F*_0_(*t*) for each ROI was computed by fitting a decaying exponential to the ROI’s time trace. When analyzing data where the stimuli contained interleaves (periods of mean gray between stimuli), only responses occurring during the interleave periods were fitted. Alternatively, with stimuli not containing interleaves, the complete time trace was fitted to calculate a baseline fluorescence. For most of the data acquired, background subtraction successfully eliminated low levels of bleed-through originating from the projector’s stimulus presentation. In cases of poor signal-to-noise recordings (particularly for the Arclight recordings and CT1 kernel extractions), custom MATLAB software used a linear model for bleed-through to subtract off contamination of the collected data. We calculated the fractional changes for each ROI trace as 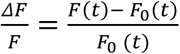.

### Imaging Data Analysis: ROI Selection

Responsive ROIs were selected as described in previous work (Matulis et al., 2020). After extracting each ROI and computing its ΔF/F, we selected desirable ROIs based on their responses to a probe stimulus—a stimulus independent of the testing stimulus that was presented at the beginning and end of each recording. For recordings of Mi1, Tm3, Mi4, and CT1, selected ROIs responded to a full-field flashes with a response of appropriate polarity, or to the white bar moving in each of the four cardinal directions. For the full-field flash stimulus probe, we selected ROIs with preferential responses to full-field ON flashes if recordings came from cells with ON-center receptive fields (*i.e*., Mi1, Tm3, Mi4, and CT1 medulla terminals). Alternatively, for cells with OFF-center receptive fields (*i.e*., CT1 lobula terminals), we selected ROIs with a preferential response to full-field OFF flashes. For ROIs selected based on their responses to a moving white bar, we based selection on the ROI’s response to a minimum of two directions of the moving bar. This white moving bar stimulus probe was used for selection in a subset of Mi1, Tm3, and Mi4 recordings.

Selection of T4 ROIs was done using procedures previously described (Salazar-Gatzimas et al., 2016, 2018). The stimulus probe for T4 recordings consisted of square waves moving right, left, up, or down, as well as light and dark edges moving rightward or leftward. The single edges section of the probe was used to determine direction selectivity indices (DSIs) and edge selectivity indices (ESIs). We then selected ROIs that met the specific response threshold previously indicated (i.e., ESI > 0.3, DSI > 0.4 for the T4 progressive layer and DSI < –0.4 for T4 regressive layer). T4 ROIs in the progressive and regressive layers were selected if they met the light vs. dark edge selectivity threshold.

### Filter extraction

For recordings of Mi1, Tm3, Mi4, and CT1, linear filters were extracted to a binary, stochastic white noise stimulus of 0.9 contrast. We used ordinary least squares (OLS) regression to compute the linear filter that best predicted neural responses. Concretely, to compute the filter, we solved the equation ***Sk*** = ***r***, where ***r*** is the response vector, and ***S*** is a matrix of stimulus contrasts that preceded each response. This used *N* pairs of stimulus-vectors and responses, as follows:

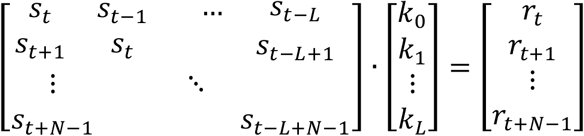

The stimulus and response values at a specific time *t* are *s_t_* and *r_t_*, respectively. The filter is *L* + 1 elements long, and *k_i_* gives the filter’s value at a specific time lag, *i*. We used standard methods in Matlab to solve this over-determined ordinary least square equation to obtain the best fit kernel ***k***. In the equations above, we included stimulus values that came after each response to obtain (acausal) kernel elements with negative lag times.

For Arclight kernels (Figure S5), we used a temporal super-resolution method that allowed us to extract the kernels with high resolution (~120 Hz) even while sampling responses at 13 frames per second (Mano et al., 2019).

As described previously, nonlinearities were computed by fitting each fly’s response to a linear-nonlinear (LN) model (Matulis et al., 2020). In this model, the binary flickering stimulus was linearly filtered by the fitted kernel and then acted on by an instantaneous nonlinearity. To plot the nonlinearities, the linear prediction was plotted against the measured responses, with individual points binned by their linear prediction to determine a non-parametric nonlinearity. This nonlinearity represents the nonlinearity associated with the transformation of visual contrast to calcium (Yang et al., 2016) and the nonlinearity associated with our calcium indicator (Chen et al., 2013). In a LN model, if only the filter amplitude changes, then the plotted nonlinearities will lie on top of one another. These nonlinearities would be expected to change if, for instance, the basal calcium level in a cell changed under a manipulation.

To deconvolve linear calcium indicator dynamics from the filter, we assumed that the indicator acted as a first order low-pass filter with time constant of 250 ms (Chen et al., 2013). We then solved the same ordinary least squares equation above, except that we modeled the response as a first-order inhomogeneous recurrence relation with variable source:

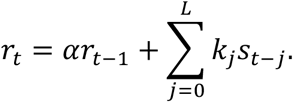

We chose the parameter 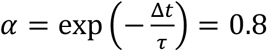, where *τ* = 250 ms is the filter time constant and Δ*t* = 70ms is our imaging measurement interval, which follows from comparing the formal solution to this recurrence to its analog in continuous time. This changed the equation above to read:

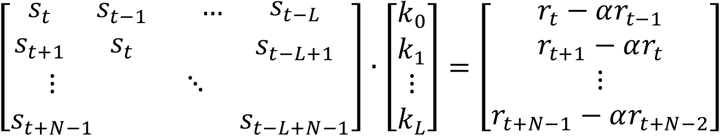

We then used the same method to solve for this deconvolved ***k***.

### Behavioral analysis

Fly optomotor turning responses were measured and quantified using methods described in previous studies (Clark et al., 2014; Creamer et al., 2019; Salazar-Gatzimas et al., 2016). Briefly, flies were temporarily anesthetized on ice, glued to metal needles using UV-cured epoxy, and tethered so they could walk on air-suspended balls. The flies were positioned in the center of panoramic screens that cover 270° of azimuth and 106° of vertical visual space. Using the monochrome green light (peak 520 nm and mean luminance of ~100 cd m^−2^) of a Lightcrafter DLP (Texas Instruments, USA), we projected stimuli onto the screens creating a virtual cylinder around the fly. Turning response was quantified by measuring the rotation of the ball at 60 Hz using an optical mouse sensor. Flies were tested in a warm, temperature-controlled behavioral chamber (34–36°C), which resulted in strong behavior. Flies were presented with a periodic, white bar velocity sweep stimulus for 1 second trials with a velocity chosen in a pseudorandom order (table S3). Turning responses were averaged over the duration of the stimulus presentation and over trials to create fly averages. These were then averaged across multiple flies. Tuning curves were created following the same analysis procedure as in T4 imaging data.

### Tuning curves and center of mass

To compute the T4 tuning curves in Figures 4–5, S9–10, and S13, we recorded T4 responses to a periodic stimulus moving at a variety of speeds. Each stimulus lasted 5 seconds. For each ROI, we computed the mean response over the 5 second presentation and over all presentations of a specific speed. We then averaged ROIs within flies to generate each fly’s tuning curve. These were averaged across flies and those average curves were presented as normalized (Figure 4–5) or not normalized (Figure S9).

These tuning curves were quantified with a single number representing the center of mass of the curve as a function of log-velocity. The center of mass was computed as:

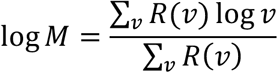

where *R*(*v*) was the mean Δ*F/F* response to a specific velocity *v*. *R*(*v*) was set to be 0 if values were negative. Thus, the center of mass is a geometric mean of the velocities, weighted by the responses.

#### Numerical modeling

##### Synaptic models for T4 neurons

We constructed synaptic models for T4 neurons following prior work (Zavatone-Veth et al., 2020). Here, we briefly summarize this synaptic model, and describe two elaborations introduced in this work. The previously-introduced model (Zavatone-Veth et al., 2020) includes three inputs: a delayed ND-offset OFF inhibitory input representing Mi9, a centered ON excitatory input representing Mi1 and Tm3, and a delayed PD-offset ON inhibitory input representing Mi4. All inputs are modeled as linear-nonlinear (LN) transformations of the input contrast. Each input has a Gaussian spatial acceptance function with a full width at half maximum of 5.7 degrees (Stavenga, 2003; Zavatone-Veth et al., 2020); we denote the spatially filtered contrast signal by *c*(*t, x*) for brevity. For temporal filters *f*_Mi9_, *f*_Mi1/Tm3_, and *f*_Mi4_, the three inputs to the model cell are then defined as rectified linear functions that mimic the polarity-selectivity of inputs to T4 cells:

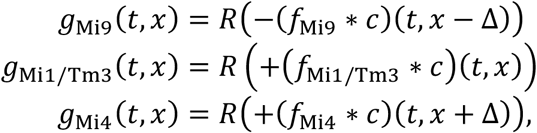

where * denotes temporal convolution, *R*(*x*) = max{0, *x*} is the ramp function, and Δ = 5° is the spacing between neighboring inputs (Stavenga, 2003; Zavatone-Veth et al., 2020). Using these inputs, we then define the conductances for excitatory and inhibitory currents:

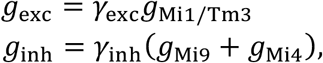

where *γ*_exc_ and *γ*_inh_ are constant gain factors. We then define the membrane potential *V*_m_ of the model T4 cell as

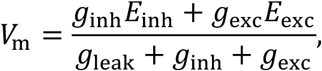

where *E*_inh_ and *E*_exc_ are the reversal potentials for inhibitory and excitatory currents, respectively, and *g*_leak_ is the leak conductance. Briefly, this nonlinearity follows from defining *V*_m_ such that the reversal potential for leak currents is 0 mV and then making a pseudo-steady-state approximation for the voltage in the limit of small membrane capacitance (Gruntman et al., 2018; Torre and Poggio, 1978; Zavatone-Veth et al., 2020). Finally, we model the transformation from membrane voltage to calcium concentration by a positively rectifying half-quadratic function *R*^2^(*x*) ≡ (*R*(*x*))^2^:

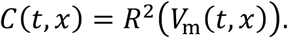

The gain factors *γ*_exc_ and *γ*_inh_ can then be represented in units of *g*_leak_; as in prior work (Zavatone-Veth et al., 2020) we fix 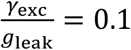 and 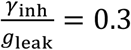 throughout. We note that this choice also reflects a choice of scale of the temporal filters; we scale all temporal filters to have unit ℓ_2_ norm after discretizing time in our simulations (Zavatone-Veth et al., 2020). This choice of scale yields filters with units of inverse contrast.

In this work, we introduce two minimally elaborated versions of this model. First, as we perform simulations using measured, non-identical temporal filters for Mi1 and Tm3, we introduce an extension with separate inputs to represent these neurons,

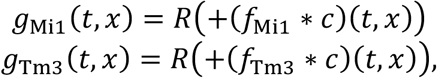

which are then integrated as

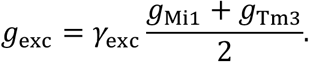

Second, we introduce a variant that incorporates a second PD-offset delay line to represent CT1, with an additional input

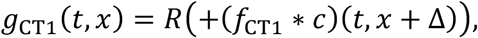

and the conductance of inhibitory currents modified to

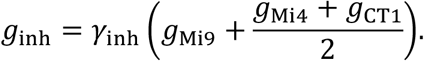

In both cases, we choose to introduce the new inputs such that the elaborated models reduce to the un-elaborated model when the relevant temporal filters are identical. We note that, with our stimulus design and chosen thresholds for the model, the Mi9-like input does not contribute to simulated model responses.

In Figure S14, we sweep the gain factor of the Mi4-like input to the model. Concretely, we fractionally rescale the value for the gain factor chosen in (Zavatone-Veth et al., 2020) by a factor ranging between zero and four. To visualize the resulting tuning changes, we plot the center of mass of each tuning curve in log-velocity space.

##### Synthetic filters

As in (Zavatone-Veth et al., 2020), we use an *L*_2_-normalized second order lowpass filter 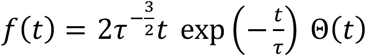 and its normalized distributional derivative 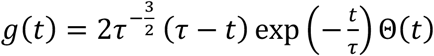, with rescaling to obtain unit ℓ_2_ norms after discretization. The function Θ(*t*) is the Heaviside step function.

##### Visual Stimuli

In all simulations, we used 5-degree-wide drifting bar stimuli with a spatial period of 45 degrees, designed to mimic the stimuli used in experiments. We chose the background of these stimuli to have contrast zero, and the foreground bars to have contrast one. Therefore, the Mi9-like input of the model from (Zavatone-Veth et al., 2020) does not respond to these stimuli, as it is sensitive only to negative contrasts.

##### Numerical methods

As in prior work (Zavatone-Veth et al., 2020), all simulations were performed using a spatial sampling interval of 0.5 degrees and a temporal sampling interval of 1/240 s. All simulations were performed using Matlab 9.8 (R2020a) (The MathWorks, Natick, MA, USA).

##### Smoothing measured temporal filters using discrete Laguerre functions

We smoothed the measured, calcium-deconvolved filters by projecting them into a truncated basis set of discrete Laguerre functions (Mano et al., 2019; Marmarelis, 1993). For a scale parameter *α* ∈ (0,1), the discrete Laguerre polynomials 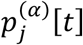 are the orthogonal polynomials on 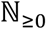 for the discrete exponential weight, i.e., the polynomials satisfying 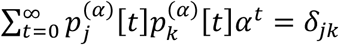. The orthonormal discrete Laguerre functions 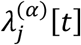 then follow by absorbing the weight, and are explicitly given as

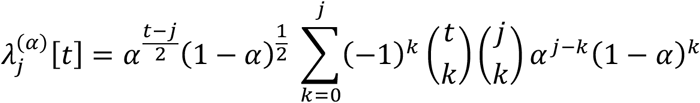

for 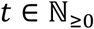 and 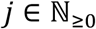. These functions form a complete orthonormal basis for the space of square-summable functions on 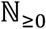, and are a convenient basis for temporal kernels as they incorporate the expected temporal decay (Marmarelis, 1993). As in prior work (Mano et al., 2019), we chose the five lowest-order functions. To obtain qualitatively reasonable smoothed filters, we set *α* = 0.2 (Marmarelis, 1993). After projecting the deconvolved filters into this subspace, we re-normalized them to have unit ℓ_2_ norm. The resulting smoothed filters are plotted along with their deconvolved and raw counterparts in Figure S11.

##### Statistical analysis

For statistical purposes, individual flies were considered independent measurements. Each fly yielded multiple ROIs, and the ROIs’ responses were averaged together to generate a single response per fly. In extracting filters, all the filters extracted from a fly’s multiple ROI traces were averaged to obtain a single filter per fly. Then, each fly’s filter was normalized by its peak amplitude, and each filter’s characteristic rise, peak, and fall time were computed. To display normalized average filters, they were averaged across flies, before the average was scaled to have a maximum excursion of 1. Similarly, the dynamics bar plots are also the average across multiple flies. The solid filter line and shaded error bars indicate the mean ± SEM. Similar averaging was done for T4 recordings. After averaging ROI traces in time, a single tuning curve was obtained for the progressive and regressive layers of T4 and T5. Main text figures depict a tuning curve resulting from the combination of the progressive and regressive layers for T4. All tuning curves were normalized to their peak on a per-fly basis and the curve’s center of mass was computed on a per fly basis. In the figure legends, n values indicate the number of individual flies. Some control genotypes were tested continuously throughout the course of experiments, which were performed over several years. This is reflected in larger sample sizes for those genotypes. Throughout, non-parametric tests were used to assess statistical significance, as noted in the figure legends.

## Acknowledgements

This paper benefitted from discussions with J. Demb, M. Higley, L. Kaczmarek, H. Keshishian, P. Masset, and S. Qin, as well as members of the Clark lab. The UAS-*slo* construct was provided by N. Atkinson. The UAS-*Sh*-DN construct was given to us by H. Keshishian.

## Funding

Support of ADG-S: Ford Foundation Fellowships Program, the NSF Graduate Research Fellowships Program, the Philanthropic Educational Organization Sisterhood Fellowship, and the Kavli Scholar Award Fellowship.

Support of JAZ-V: NSF-Simons Center for Mathematical and Statistical Analysis of Biology at Harvard and the Harvard Quantitative Biology Initiative.

Support of DAC and this project: NIH R01EY026555, NIH R01NS121773, NIH P30EY026878, NSF IOS1558103, the E. Mathilda Ziegler Foundation for the Blind, and a Sloan Foundation Research Fellowship in Neuroscience.

## Author Contributions

Conceptualization: ADG-S and DAC.

Methodology & Investigation: ADG-S, CAM, and BAB

Analysis: ADG-S, JAZ-V, JC, CAM, and DAC.

Modeling: ADG-S, JAZ-V, and DAC.

Writing: ADG-S and DAC.

## Data and Code Availability

Data is available upon request to the Lead Contact, Damon Clark. Software for the models is available from GitHub: http://www.github.com/ClarkLabCode/SynapticModelTimingCode.

## Supplementary Materials

### Supplementary Text

**Figure S1.**
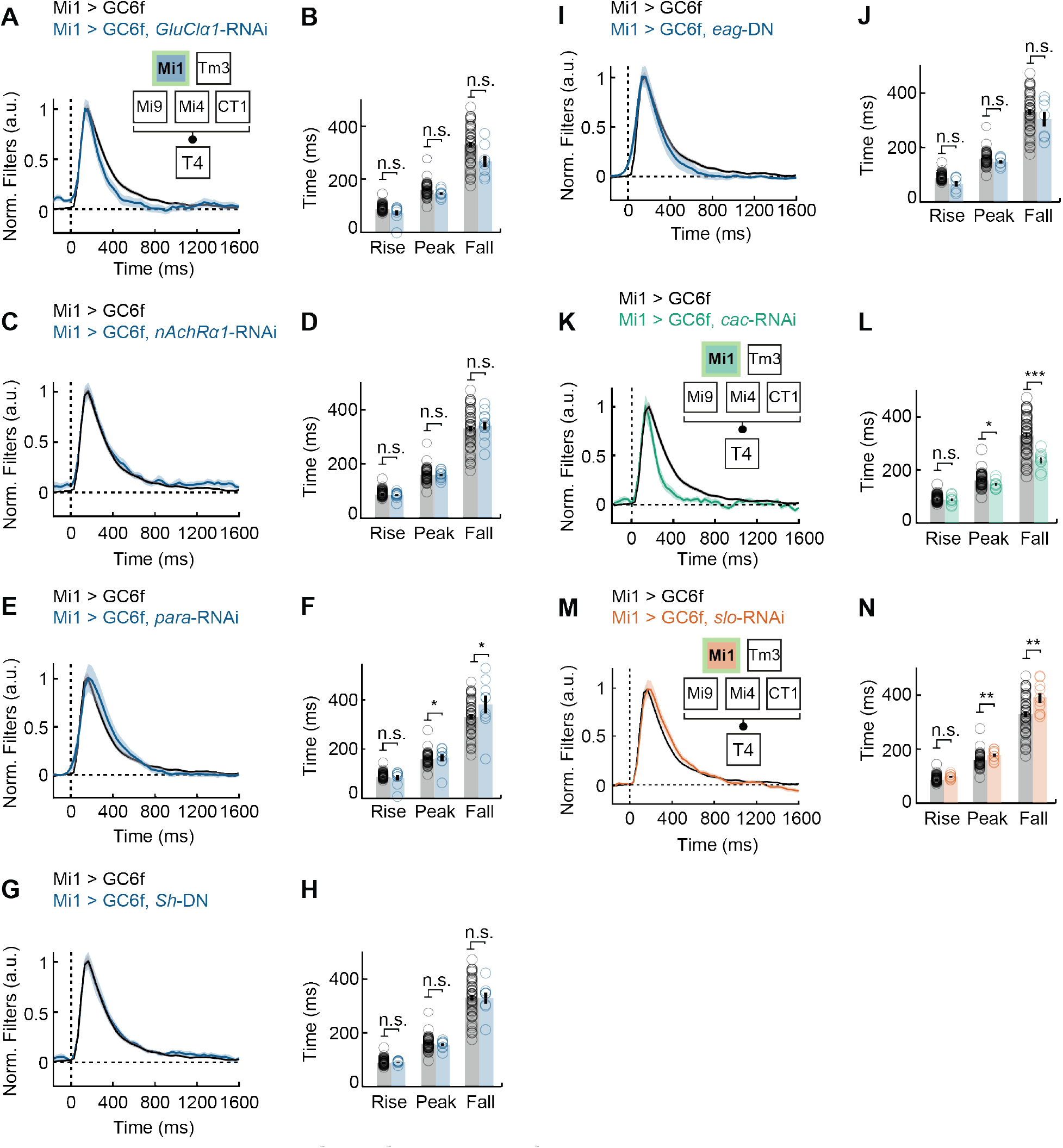
Suppression of *slowpoke* and *cacophony* expression significantly alters Mi1 filter dynamics. **(A)** Filters of Mi1 with the glutamate-gated Cl^−^ channel *GluClα1* knocked-down (Mi1 > GC6f, *GluClα1*-RNAi, n = 9), compared to wildtype Mi1 (Mi1 > GC6f, n = 68). Lines are mean ± SEM. **(B)** Filter dynamics quantification of (A): filter’s half-rise (rise), peak, and half-fall (fall) times averaged across flies. **(C)** As in (A), but for Mi1 with the nicotinic acetylcholine α1 receptor knocked-down (Mi1 > GC6f, *nAchRα1*-RNAi, n = 12), compared to wildtype Mi1 (Mi1 > GC6f, n = 68). **(D)** As in (B), but for filters in (C). **(E)** As in (A), but for Mi1 with the voltage-gated Na^+^ channel *para* knocked-down (Mi1 > GC6f, *para*-RNAi, n = 9), compared to wildtype Mi1 (Mi1 > GC6f, n = 68). **(F)** As in (B), but for filters in (E). **(G)** As in (A), but for Mi1 with a dominant negative mutation in the voltage-gated K^+^ channel *Shaker* (Mi1 > GC6f, *Sh*-DN, n = 8), compared to wildtype Mi1 (Mi1 > GC6f, n = 68). **(H)** As in (B), but for filters in (G). **(I)** As in (A), but for Mi1 with a dominant negative mutation in the voltage-gated delayed rectifier K^+^ channel *Ether-a-go-go* (Mi1 > GC6f, *eαg*-DN, n = 7), compared to wildtype Mi1 (Mi1 > GC6f, n = 68). **(J)** As in (B), but for filters in (I). **(K)** As in (A), but for Mi1 with the voltage-gated Ca^2+^ channel *cacophony* (*cac*), knocked-down (Mi1 > GC6f, *cac*-RNAi, n = 9), compared to wildtype Mi1 (Mi1 > GC6f, n = 68). **(L)** As in (B), but for filters in (K). **(M)** As in (A), but for Mi1 with the voltage- and calcium-gated K^+^ channel *slowpoke* (*slo*), knocked-down (Mi1 > GC6f, *slo*-RNAi, n = 19), compared to wildtype Mi1 (Mi1 > GC6f, n = 68). **(N)** As in (B), but for filters in (M). Note that in cases where genetic manipulations did not elicit an observable phenotype, we do not interpret the absence of a change as indicating that the gene is not necessary for wildtype dynamics, since there are a host of reasons why such experiments could have failed to show a phenotype. (* p<0.05, ** p<0.01, *** p<0.001 by Wilcoxon signed-rank tests across flies.)

**Figure S2.**
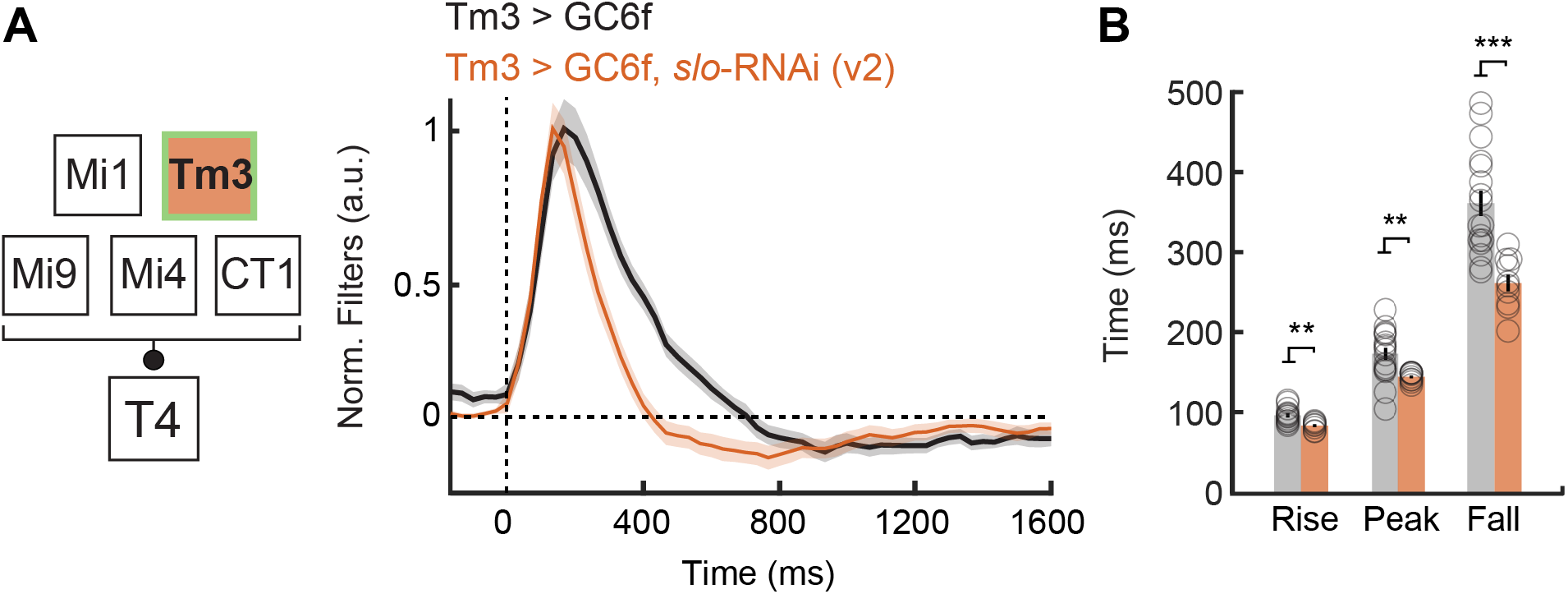
Independent *slowpoke* knock-down in Tm3 speeds up the cell’s dynamics. **(A)** Filters of Tm3 with the Ca^2+^-gated K^+^ channel *slowpoke* (*slo*) knocked-down (*slo*-RNAi) (Tm3 > GC6f, *slo*-RNAi (v2), n = 10) compared to wildtype Tm3 (Tm3 > GC6f, n = 17). This *slo*-RNAi construct was obtained from an independent RNAi library (VDRC, labeled v2 here) (Dietzl et al., 2007). Lines are mean ± SEM. **(B)** Filter dynamics quantification of (A): filter’s half-rise (rise), peak, and half-fall (fall) times averaged across flies. A single outlying fly of genotype Tm3 > GC6f was removed from the analysis of fall times, since its fall time was computed to be ~1500 ms. This did not affect the significance of the difference shown. (* p<0.05, ** p<0.01, *** p<0.001 by Wilcoxon signed-rank tests across flies.)

**Figure S3.**
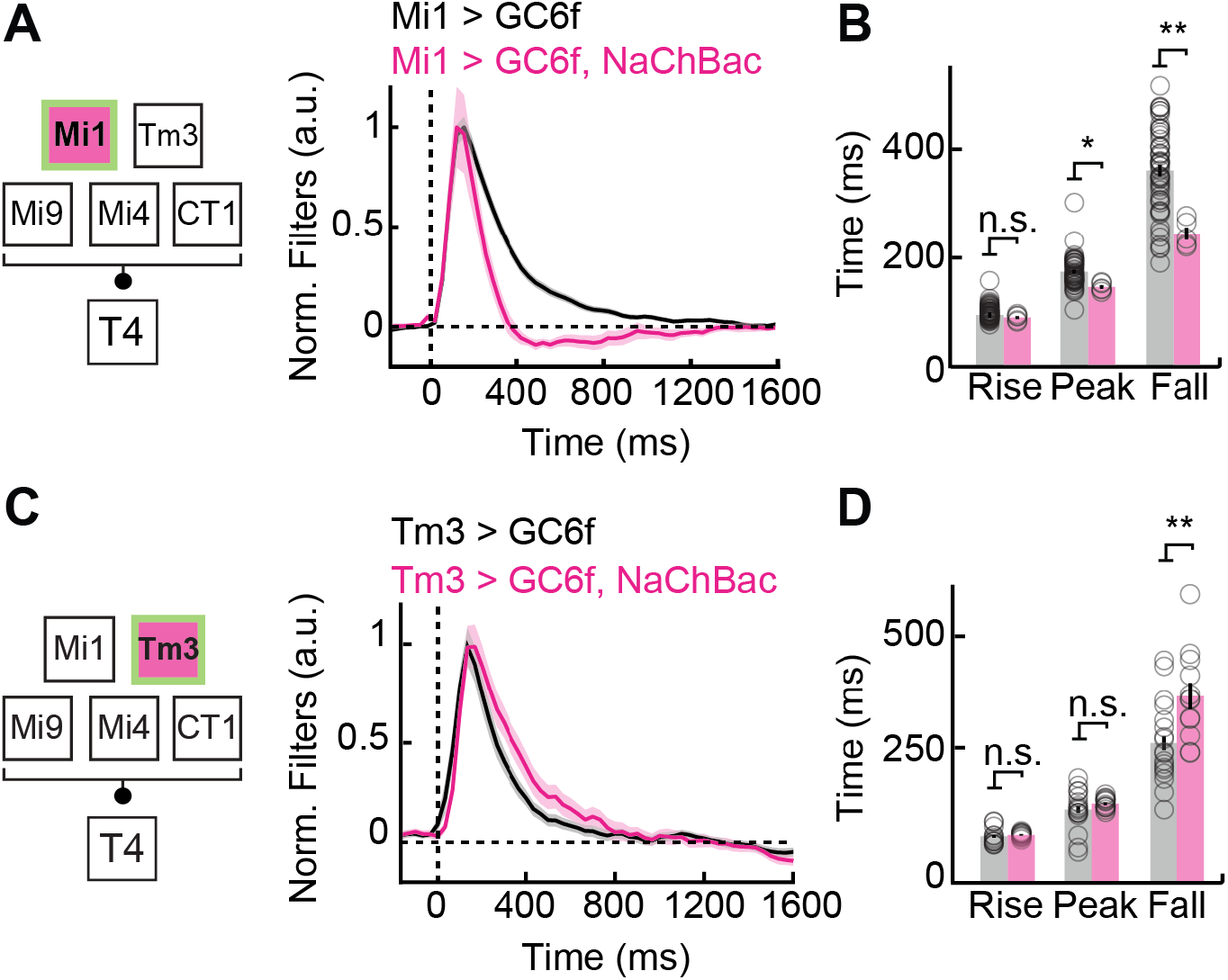
Expression of NaChBac speeds up Mi1 dynamics, but slows down Tm3 dynamics. **(A)** Filters of Mi1 expressing the bacterial, voltage-gated Na^+^ channel NaChBac (Mi1 > GC6f, NaChBac, n = 5), compared to wildtype Mi1 (Mi1 > GC6f, n = 68). Lines are mean ± SEM. **(B)** Filter dynamics quantification of (A): filter’s half-rise (rise), peak, and half-fall (fall) times averaged across flies. **(C)** As in (A), but for Tm3 expressing NaChBac (Tm3 > GC6f, NaChBac, n = 15), compared to wildtype Tm3 (Tm3 > GC6f, n = 25). **(D)** As in (B), but for filters in (C). (* p<0.05, ** p<0.01, *** p<0.001 by Wilcoxon signed-rank tests across flies.)

**Figure S4.**
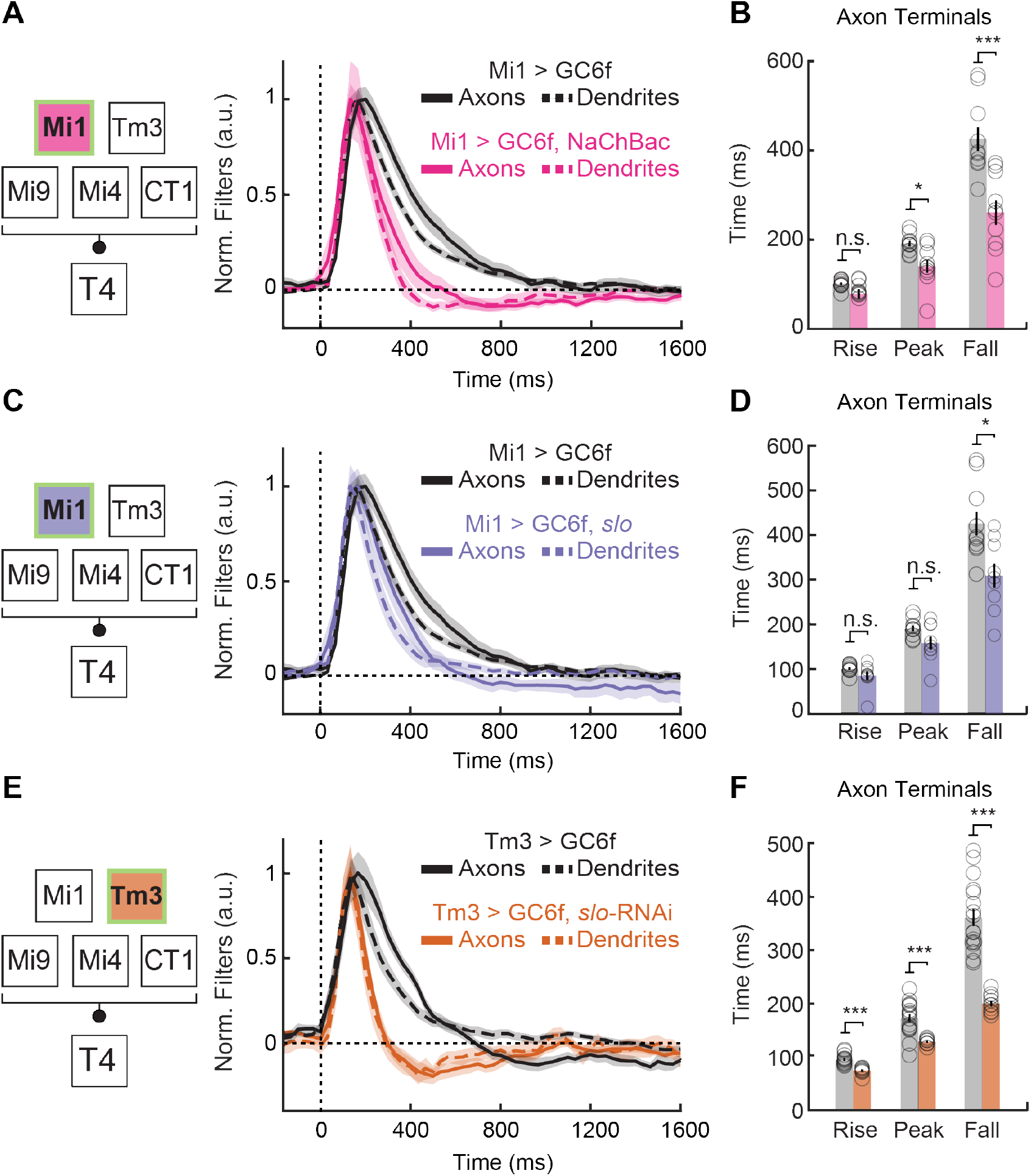
Genetic perturbations of Mi1 and Tm3 affect dendrite and axon dynamics similarly. **(A)** Dendrite filters of Mi1 expressing the bacterial, voltage-gated Na^+^ channel NaChBac (Mi1 > GC6f, NaChBac, n = 7), and axons filters of Mi1 expressing NaChBac (Mi1 > GC6f, NaChBac, n = 10), compared to wildtype Mi1 dendrites (Mi1 > GC6f, n = 68) and axons (Mi1 > GC6f, n = 10). Lines are mean ± SEM. **(B)** Axon filter dynamics quantification of (A): filter’s half-rise (rise), peak, and half-fall (fall) times averaged across flies. **(C)** As in (A), but for Mi1 over-expressing the Ca^2+^-gated K^+^ channel *slowpoke* (*slo*), (dendrites: Mi1 > GC6f, *slo*, n = 16; axons: Mi1 > GC6f, *slo*, n = 11), compared to wildtype Mi1 (dendrites: Mi1 > GC6f, n = 68; axons: Mi1 > GC6f, n = 10). **(D)** As in (B), but for filters displayed in (C). **(E)** As in (A), but for Tm3 with *slo* knock-down (dendrites: Tm3 > GC6f, *slo*-RNAi, n = 19; axons: Tm3 > GC6f, *slo*-RNAi, n = 11), compared to wildtype Tm3 (dendrites: Tm3 > GC6f, n = 25; axons: Tm3 > GC6f, n = 17). A single outlying fly of genotype Tm3 > GC6f was removed from the analysis of fall times, since its fall time was computed to be ~1500 ms. This did not affect the significance of the difference shown. **(F)** As in (B), but for filters displayed in (E). (* p<0.05, ** p<0.01, *** p<0.001 by Wilcoxon signed-rank tests across flies.)

**Figure S5.**
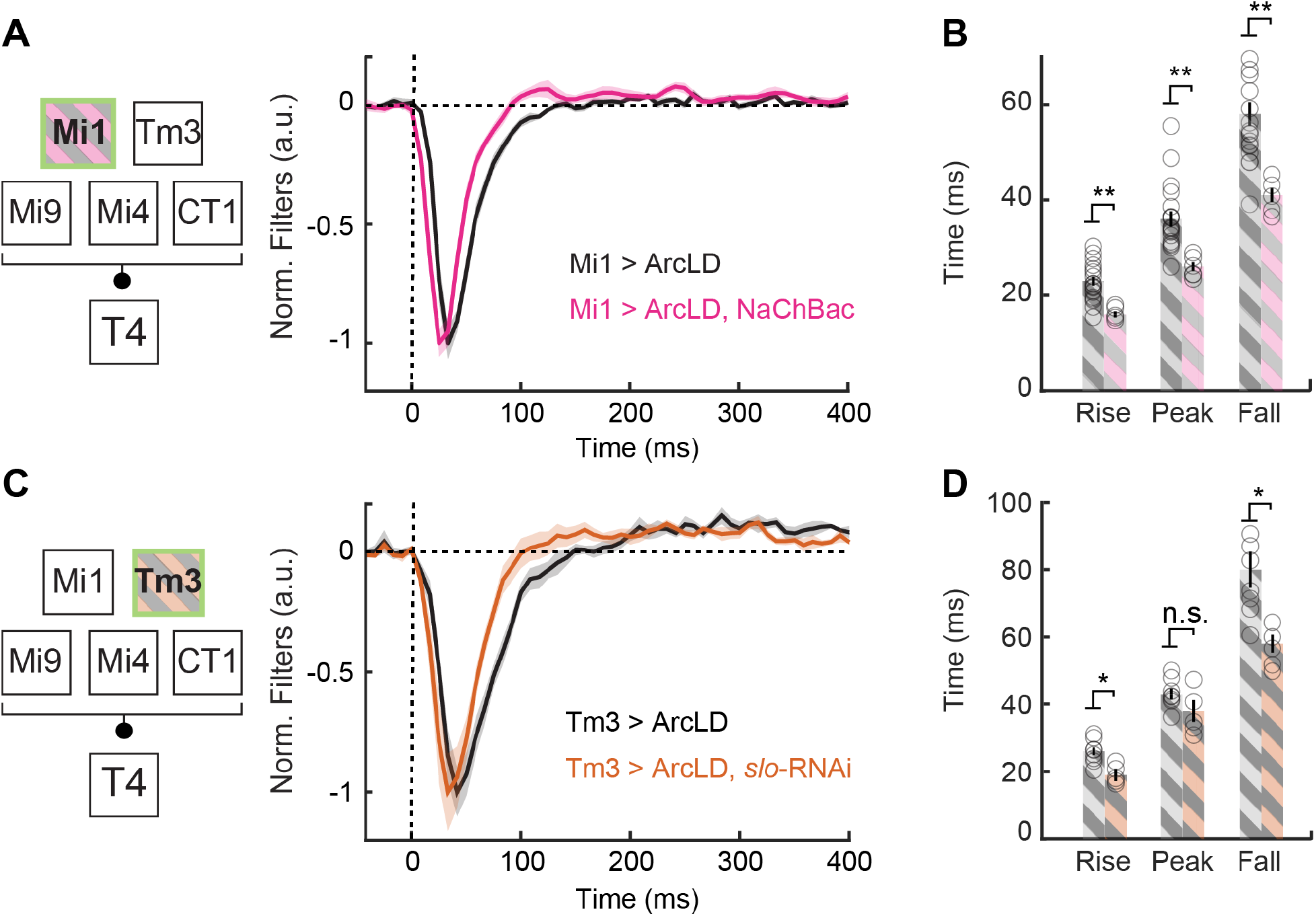
Expressing NaChBac and over-expressing *slowpoke* speeds up Mi1 and Tm3 membrane potential response dynamics. **(A)** Voltage filters of Mi1 expressing the bacterial, voltage-gated Na^+^ channel NaChBac (Mi1 > ArcLD, NaChBac, n = 5), compared to wildtype Mi1 (Mi1 > ArcLD, n = 19). Lines are mean ± SEM. Note the timescale differences from calcium filters. ArcLight fluoresces less at depolarized membrane potentials. **(B)** Filter dynamics quantification of (A): filter’s half-rise (rise), peak (max), and half-fall (fall) averaged across flies. **(C)** As in (A), but for Tm3 expressing an RNAi to knock-down the Ca^2+^-gated K^+^ channel *slowpoke* (*slo*) (Tm3 > ArcLD, *slo*-RNAi, n = 5), compared to wildtype Tm3 (Tm3 > ArcLD, n = 8). **(D)** As in (B), but for filters in (C). (* p<0.05, ** p<0.01, *** p<0.001 by Wilcoxon signed-rank tests across flies.)

**Figure S6.**
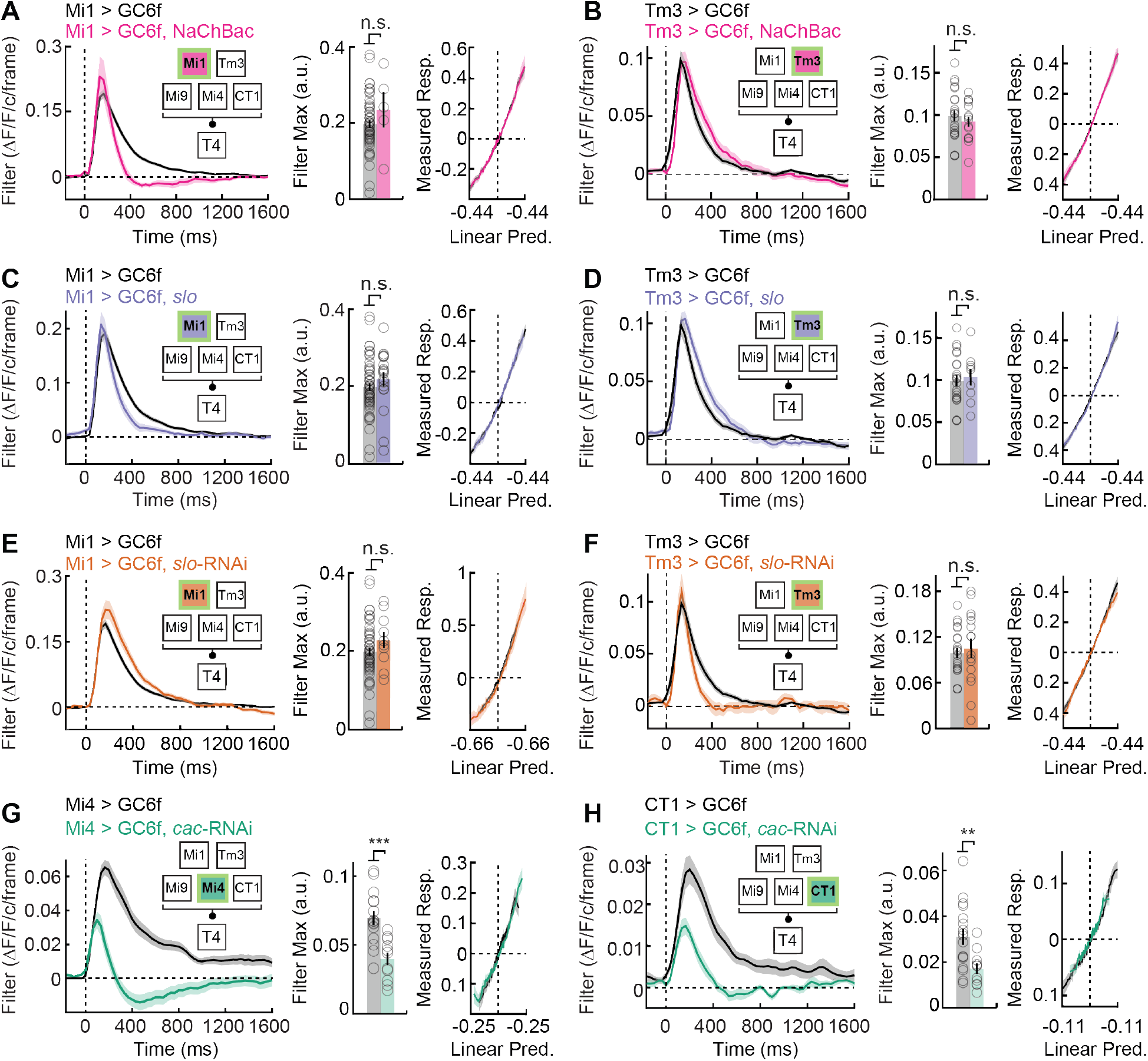
Genetic perturbations of Mi1, Tm3, Mi4, and CT1 did not strongly alter nonlinear transformations, but altered Mi4 and CT1 filter amplitude. **(A)** *Left*: un-normalized Mi1 filters expressing the bacterial voltage-gated Na^+^ channel NaChBac (Mi1 > GC6f, NaChBac, n = 7), compared to wildtype Mi1 (Mi1 > GC6f, n = 68). (Lines are mean ± SEM). Unit frames are defined as 1/30 of a second (see Methods); *Middle*: quantified maximum amplitude for filters in (A, left), on a per fly bases; *Right*: extracted nonlinearities for filters in (A, right) are based on the measured response and the linear prediction with normalized variance (see Methods). **(B)** As in (A), but for Tm3 expressing NaChBac (Tm3 > GC6f, NaChBac, n = 15), compared to wildtype Tm3 (Tm3 > GC6f, n = 25). **(C)** As in (A), but for Mi1 over-expressing the Ca^2+^-gated K^+^ channel *slowpoke* (*slo*) (Mi1 > GC6f, *slo*, n = 16), compared to wildtype Mi1 (Mi1 > GC6f, n = 68). **(D)** As in (B), but for Tm3 over-expressing *slo* (Tm3 > GC6f, *slo*, n = 8), compared to wildtype Tm3 (Tm3 > GC6f, n = 25). **(E)** As in (A), but for Mi1 with *slo* knocked-down (Mi1 > GC6f, *slo* RNAi, n = 19), compared to wildtype Mi1(Mi1 > GC6f, n = 68). **(F)** As in (B), but for Tm3 with *slo* knocked-down (Tm3 > GC6f, *slo* RNAi, n = 19), compared to wildtype Tm3 (Tm3 > GC6f, n = 25). **(G)** As in (A), but for Mi4 with the voltage-gated Ca^2+^ channel *cacophony* (*cac*) knocked-down in Mi4 (Mi4 > GC6f, *cac* RNAi, n = 11), compared to wildtype Mi4 (Mi4 > GC6f, n = 15). **(H)** As in (G), but for CT1 with *cac* knocked-down (CT1 > GC6f, *cac* RNAi, n = 11), compared to wildtype CT1 (CT1 > GC6f, n = 17). (* p<0.05, ** p<0.01, *** p<0.001 by Wilcoxon signed-rank tests across flies.)

**Figure S7.**
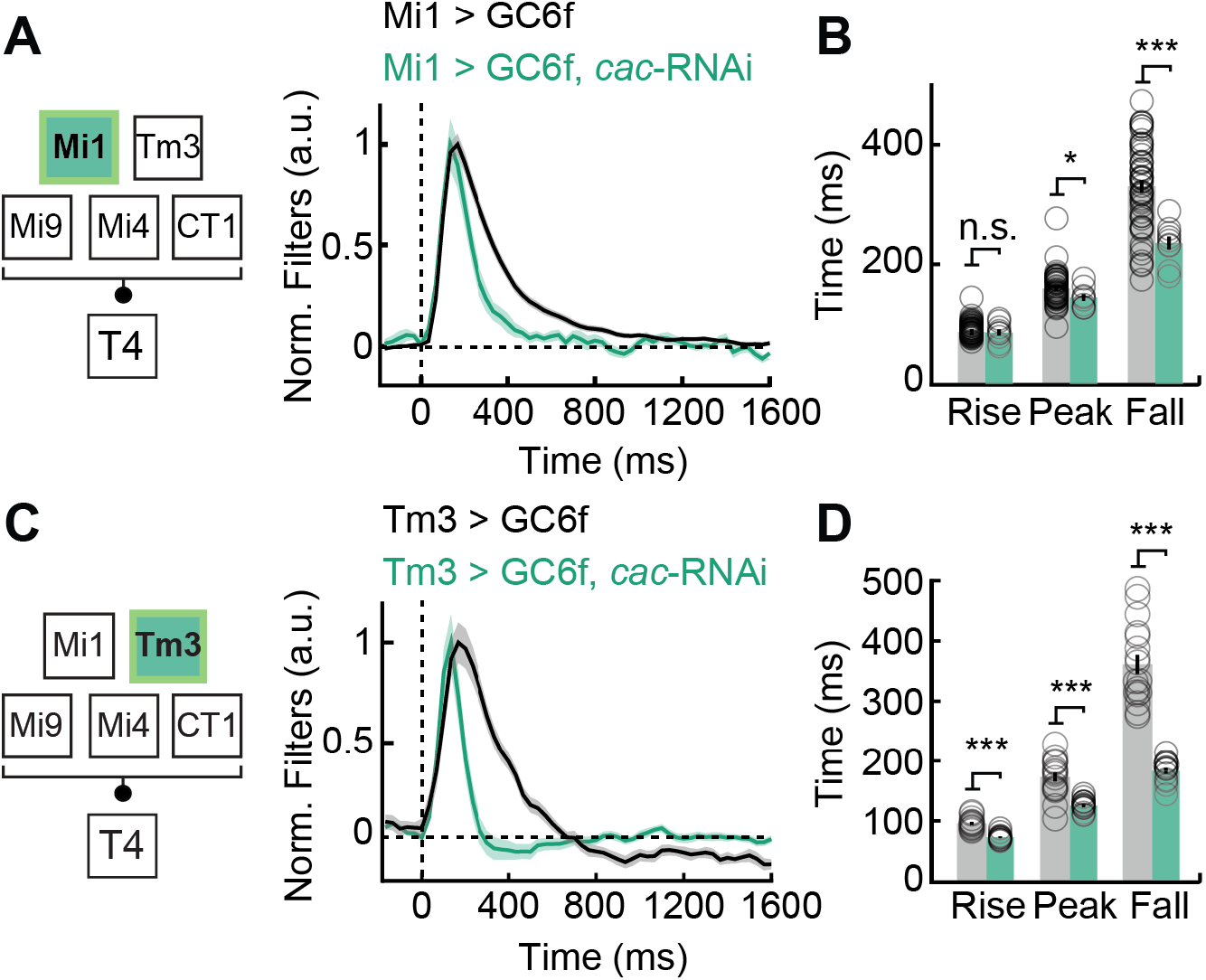
Knocking-down *cacophony* in Mi1 and Tm3 speeds up filter dynamics in both. **(A)** Filters of Mi1 with the voltage-gated Ca^2+^ channel *cacophony* (*cac*), knocked-down (Mi1 > GC6f, *cac*-RNAi, n = 9), compared to wildtype Mi1 (Mi1 > GC6f, n = 68). Lines are mean ± SEM. **(B)** Filter dynamics quantification of (A): filter’s half-rise (rise), peak, and half-fall (fall) times averaged across flies. **(C)** As in (A), but for Tm3 expressing *cac*-RNAi (Tm3 > GC6f, *cac*-RNAi, n = 10), compared to wildtype Tm3 (Tm3 > GC6f, n = 17). **(D)** As in (B), but for filters in (C). (* p<0.05, ** p<0.01, *** p<0.001 by Wilcoxon signed-rank tests across flies.)

**Figure S8.**
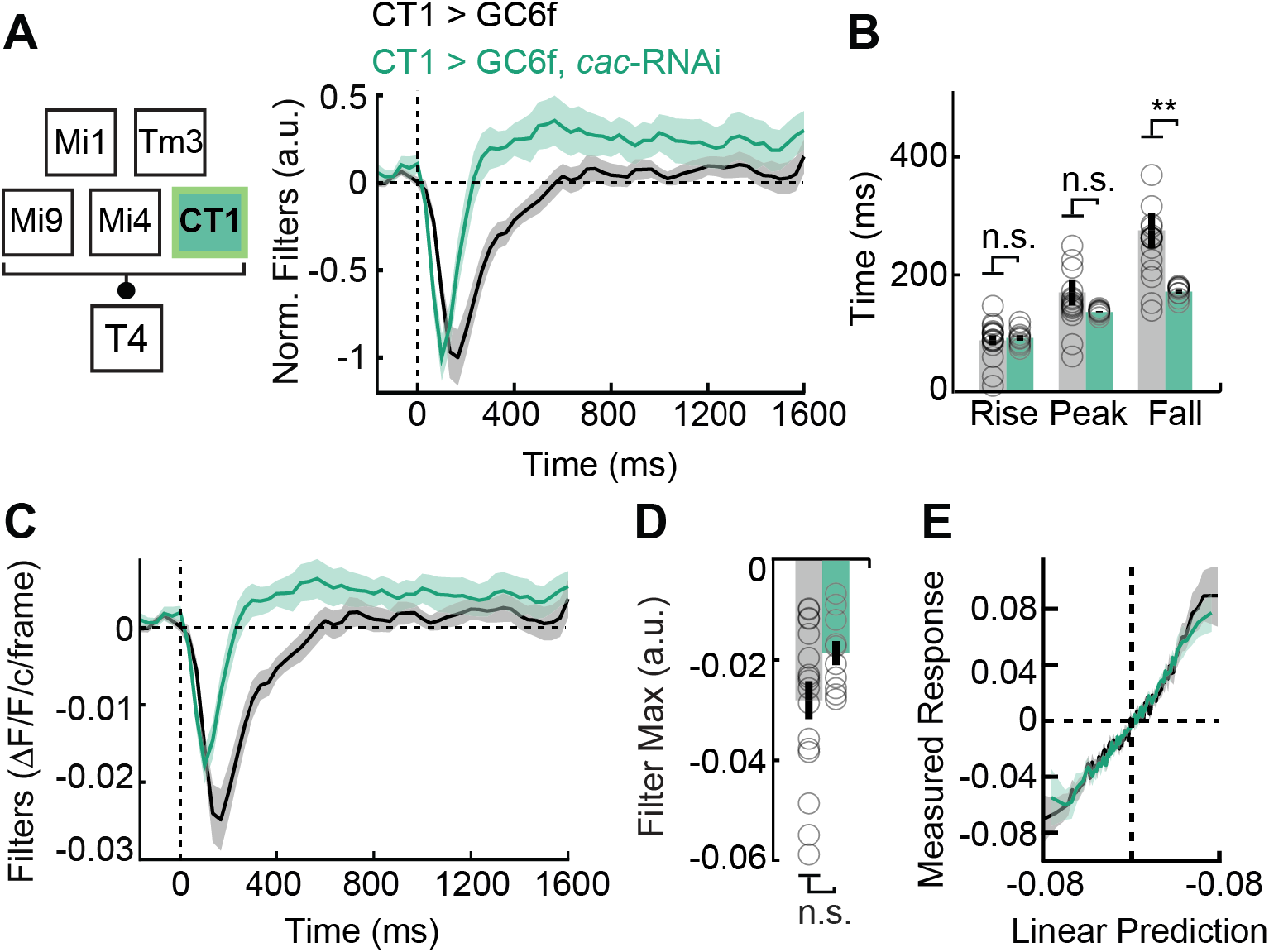
Knocking-down *cacophony* in CT1 speeds up the filter dynamics in its lobula axon terminals. **(A)** Filters of CT1 with the voltage-gated Ca^2+^ channel *cacophony* (*cac*) knocked-down (CT1 > GC6f, *cac*-RNAi, n = 10), compared to wildtype CT1 (CT1 > GC6f, n = 17). Lines are mean ± SEM. **(B)** Filter dynamics quantification of (A): filter’s half-rise (rise), peak, and half-fall (fall) times averaged across flies. **(C)** Un-normalized filters of CT1 expressing *cac*-RNAi expression (CT1 > GC6f, *cac*-RNAi, n = 10), compared to wildtype CT1 (CT1 > GC6f, n = 17). Unit frames are defined as 1/30 of a second (see Methods). **(D)** Quantified maximum amplitude for filters in (C), on a per fly bases. **(E)** Extracted nonlinearities based on measured responses and linear prediction for CT1 expressing *cac*-RNAi (CT1 > GC6f, *cac*-RNAi, n = 10), compared to wildtype CT1 (CT1 > GC6f, n = 17) (see Methods). (* p<0.05, ** p<0.01, *** p<0.001 by Wilcoxon signed-rank tests across flies.)

**Figure S9.**
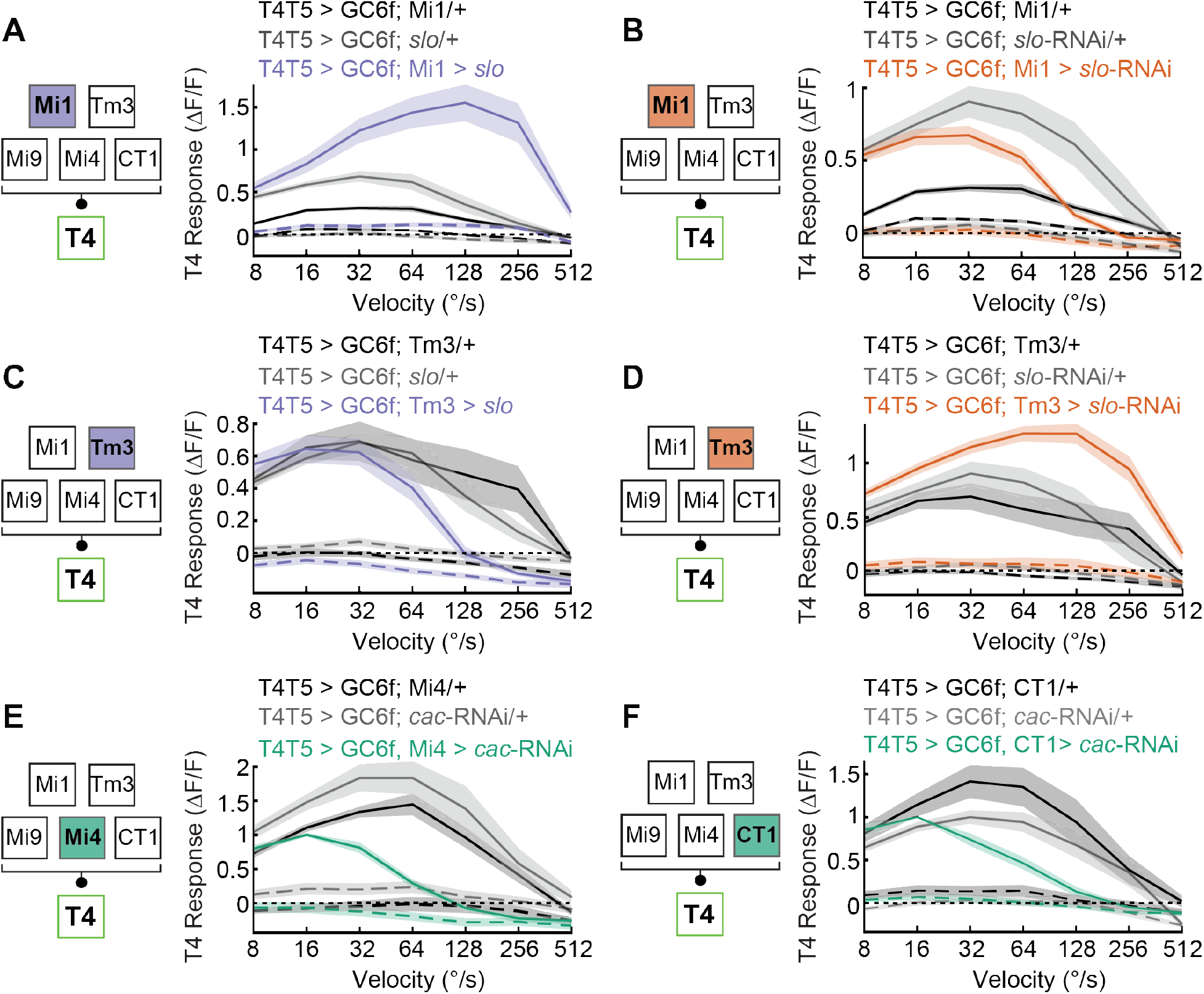
T4 tuning curves to white bars moving in the preferred and null direction. **(A)** T4 responses to white bars moving in the preferred direction (*solid lines*) and the null direction (*dashed lines*) for flies over-expressing the Ca^2+^-gated K^+^ channel *slowpoke* (*slo*) (T4T5 > GC6f, Mi1 > *slo*, n = 9) compared to two genetic controls (T4T5 > GC6f; Mi1/+, n = 43 and T4T5 > GC6f; *slo/+*, n = 11). Lines are mean ± SEM. **(B)** As in (A), but for Mi1 expressing *slo*-RNAi (T4T5 > GC6f, Mi1 > *slo* RNAi, n = 7), compared to two genetic controls (T4T5 > GC6f; Mi1/+, n = 43 and T4T5 > GC6f; *slo* RNAi/+, n = 10). **(C)** As in (A), but for Tm3 over-expressing *slo* (T4T5 > GC6f, Tm3 > *slo*, n = 7), compared to two genetic controls (T4T5 > GC6f; Tm3/+, n = 11 and T4T5 > GC6f; *slo*/+, n = 11). **(D)** As in (A), but for Tm3 expressing *slo*-RNAi (T4T5 > GC6f, Tm3 > *slo*-RNAi, n = 12), compared to two genetic controls (T4T5 > GC6f; Tm3/+, n = 11 and T4T5 > GC6f; *slo*-RNAi/+, n = 10). **(E)** As in (A), but for Mi4 expressing RNAi to knock-down the voltage-gated Ca^2+^ channel *cacophony* (*cac*) (T4T5 > GC6f, Mi4 > *cac*-RNAi, n = 8) compared to two genetic controls (T4T5 > GC6f; Mi4/+, n = 12 and T4T5 > GC6f; *cac*-RNAi/+, n = 8). **(F)** As in (A), but for CT1 expressing *cac*-RNAi (T4T5 > GC6f, CT1 > *cac*-RNAi, n = 7) compared to two genetic controls: T4T5 > GC6f; CT1/+ (n = 12) and T4T5 > GC6f; *cac*-RNAi/+ (n = 8). (* p<0.05, ** p<0.01, *** p<0.001 by Wilcoxon signed-rank tests across flies.)

**Figure S10.**
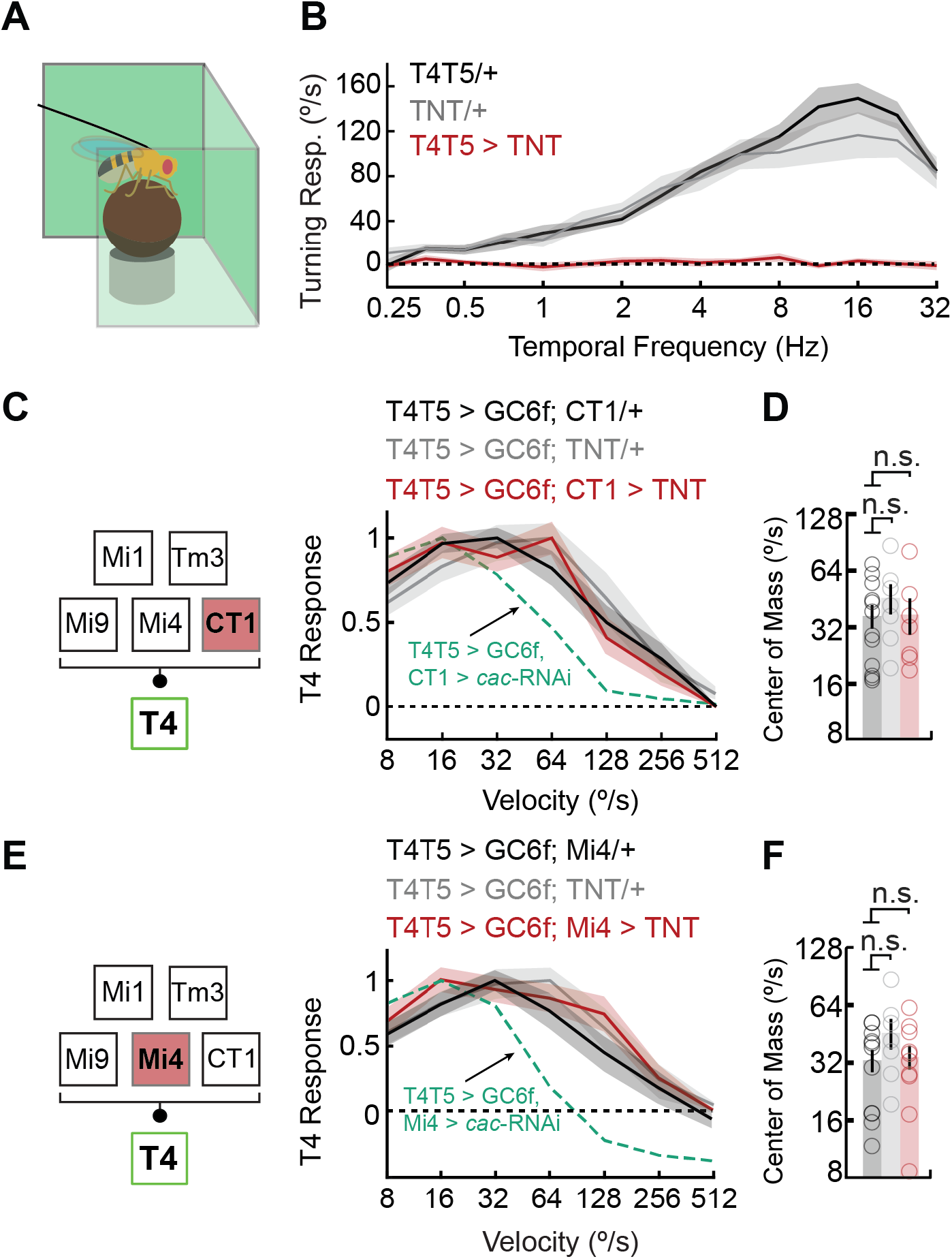
Silencing CT1 or Mi4 with tetanus toxin does not affect T4 tuning. **(A)** A fly-on-the-ball setup was used to measure flies’ behavioral turning response. **(B)** Flies’ turning responses to sinewave gratings of various temporal frequencies were recorded. Flies expressing tetanus toxin (TNT) in T4 and T5 (T4T5 > TNT, n = 11) were compared to two genetic controls (T4T5/+, n = 15 and TNT/+, n = 13). **(C)** T4 tuning curves of flies expressing TNT in CT1 (T4T5 > GC6f, CT1 > TNT, n = 7), compared to two genetic controls (T4T5 > GC6f; CT1/+, n = 12 and T4T5 > GC6f; TNT/+, n = 7). Dashed line represents the tuning curve of T4T5 > GC6f, CT1 > *cac*-RNAi. Lines are mean ± SEM. **(D)** The tuning curve’s center of mass is a weighted average of each tuning curve shown in (C), plotted in log-velocity space. **(E)** As in (C), but for Mi4 expressing TNT (T4T5 > GC6f, Mi4 > TNT, n = 10), compared to two genetic controls (T4T5 > GC6f; Mi4/+, n = 12 and T4T5 > GC6f; TNT/+, n = 7). Dashed line represents the tuning curve of T4T5 > GC6f, Mi4 > *cac*-RNAi. **(F)** As in (D), but for tuning curves shown in (E). (* p<0.05, ** p<0.01, *** p<0.001 by Wilcoxon signed-rank tests across flies.)

**Figure S11.**
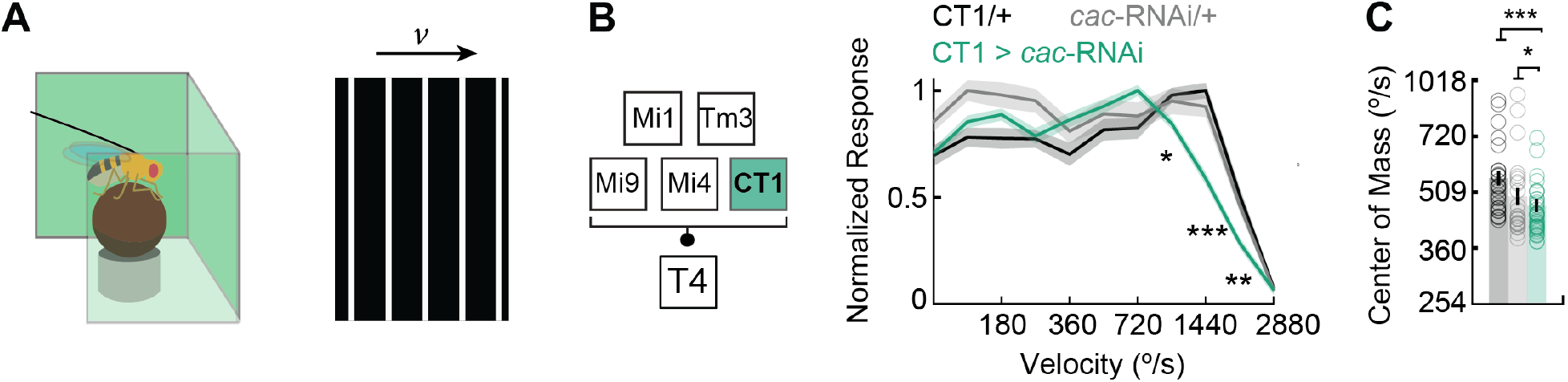
Knock-down of *cacophony* in CT1 mediates the dynamics of flies’ turning responses. **(A)** A fly-on-the-ball setup (*left*) was used to measure flies’ behavioral turning response to moving periodic, white bars at various velocities (*right*). **(B)** Flies expressing an RNAi to knock-down the voltage-gated Ca^2+^ channel *cacophony* (*cac*) in CT1 (CT1 > *cac*-RNAi, n = 40), compared to two genetic controls (CT1/+, n = 27 and *cac*-RNAi/+, n = 27). Lines are mean ± SEM. (* p<0.05, ** p<0.01, *** p<0.001 by Wilcoxon signed-rank tests across flies.) The difference in the velocity scale between behavioral responses and T4 and T5 measurements has been well-documented (Creamer et al., 2018; Strother et al., 2017). **(C)** The tuning curve’s center of mass is a weighted average of each tuning curve shown in (B), plotted in log-velocity space. (* p<0.05, ** p<0.01, *** p<0.001 by Wilcoxon signed-rank one-tail tests across flies.)

**Figure S12.**
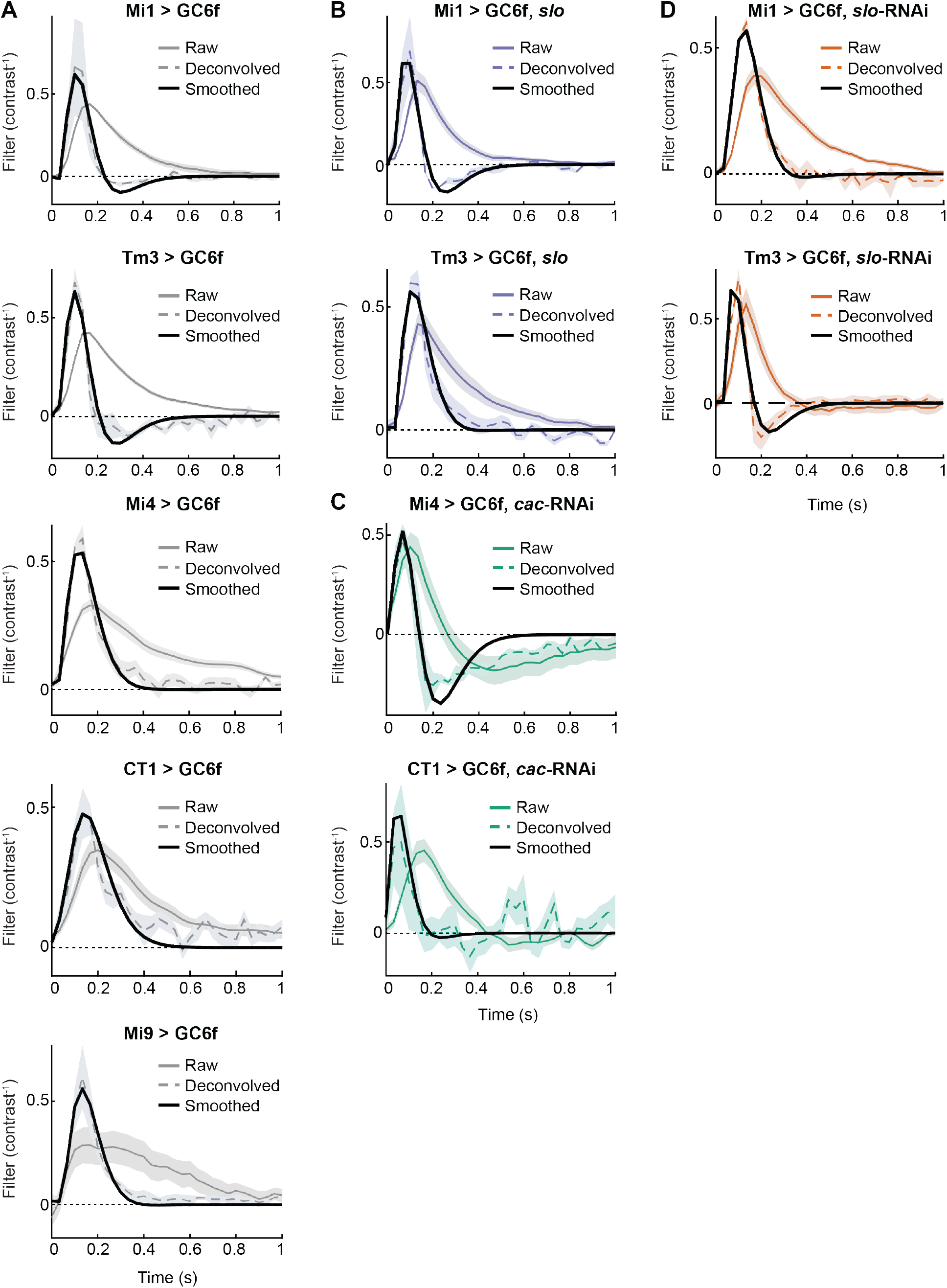
Raw, de-convolved, and smoothed filters used in synaptic model. **(A)** Wildtype filters, filters with de-convolved indicator dynamics (see Methods), and smoothed filters (see Methods) from wildtype Mi1 (Mi1 > GC6f), Tm3 (Tm3 > GC6f), Mi4 (Mi4 > GC6f), CT1 (CT1 > GC6f), and Mi9 (Mi9 > GC6f) (from top to bottom). **(B)** As in (A), but for Mi1 and Tm3 over-expressing the Ca^2+^-gated K^+^ channel *slowpoke* (*slo*) (Mi1 > GC6f, *slo* and Tm3 > GC6f, *slo*) (from top to bottom). **(C)** As in (A), but for Mi4 and CT1 expressing RNAi to knock-down the voltage-gated Ca^2+^ channel *cacophony* (*cac*) (Mi4 > GC6f, *cac*-RNAi and CT1 > GC6f, *cac*-RNAi) (from top to bottom). **(D)** As in (A), but for Mi1 and Tm3 expressing RNAi to knock-down *slo* (Mi1 > GC6f, *slo*-RNAi and Tm3 > GC6f, *slo*-RNAi) (from top to bottom).

**Figure S13.**
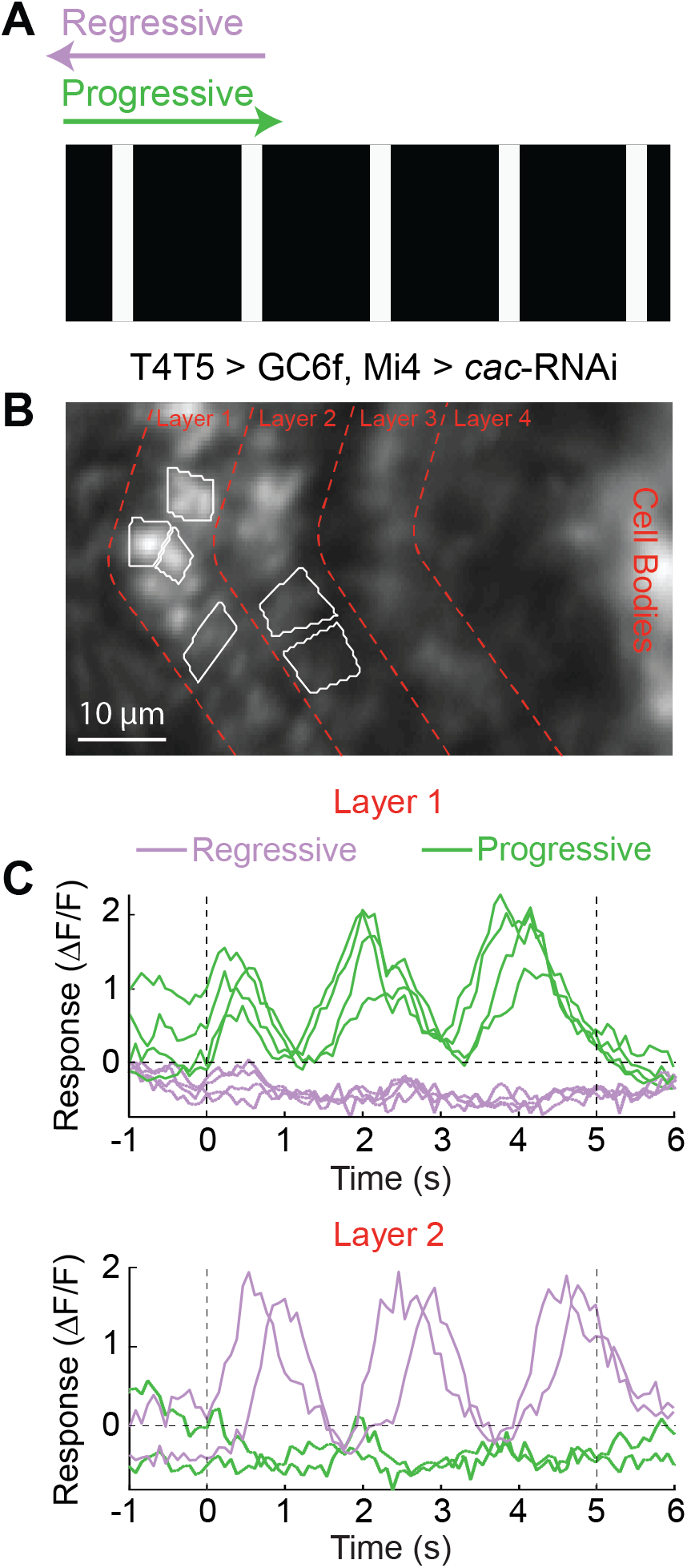
*Cacophony* knock-down in Mi4 does not switch the directionality of the progressive and regressive layers in the lobula plate. **(A)** White 5°-wide bars, with 30° spacing rotate in progressive (front-to-back) and regressive directions (back-to-front) over the eye at several velocities (8-512°/s). **(B)** Regions of interest (ROIs) are selected for two of the four anatomically-restricted layers of T4 axons in a mean two-photon microscopy image of flies where the voltage-gated Ca^2+^ channel *cacophony* (*cac*), is knocked-down (T4T5 > GC6f, Mi4 > *cac*-RNAi). ROIs in layer 1 respond to progressive stimuli (ROIs n = 4, *green*), while ROIs in layer 2 respond to regressive stimuli (ROIs n = 2, *purple*). To discriminate between ON-responding and OFF-responding ROIs, an edge selectivity index was computed from responses to light and dark edges (see Methods). **(C)** Raw change in fluorescence of T4 ROIs responding to white bars rotating at 16 °/s. ROIs selected in layer 1 (*top panel*) versus those selected in layer 2 (*bottom panel*). Green lines correspond to stimuli moving in the regressive direction, while purple lines correspond to stimuli moving in the progressive direction.

**Figure S14.**
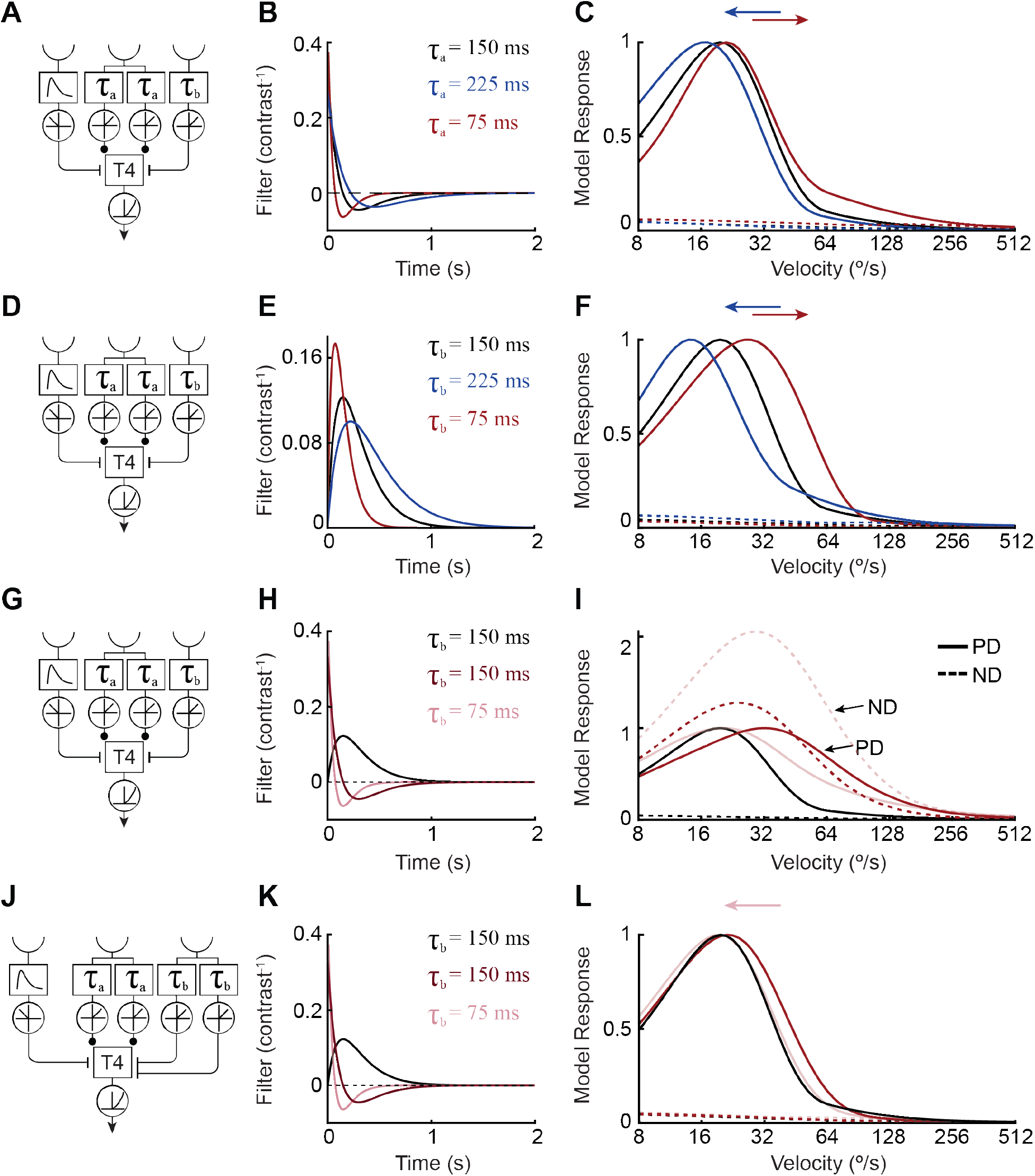
Three-input synaptic model responds consistently to changes in the dynamics of generated, synthetic low-pass and high-pass filters. **(A)** Manipulating the dynamics of the central, excitatory input to the synaptic model. Note that only one of the arms from this central, excitatory input was manipulated at a time; the time constant of the other arm was kept fixed at its ‘default’ value. As the two arms are otherwise identical, the results of these manipulations apply to either arm. **(B)** Synthetic ℓ*_2_*-normalized high-pass filters with standard dynamics (*τ*_a_ = 150 ms) were generated and compared to filters with slower (*τ*_a_ = 225 ms) or faster (*τ*_a_ = 75 ms) dynamics (see Methods). **(C)** Model responses to periodic white bar stimuli (see Methods) as a function of velocity for each filter set. Slowing the filter dynamics (*τ*_a_ = 225 ms) shifted the model’s response toward lower velocities. Conversely, models with faster filters (*τ*_a_ = 75 ms) preferred higher velocities. **(D)** As in (A), but for manipulations of the PD-offset ON inhibitory input (*τ*_b_). **(E)** As in (B), but for the ℓ*_2_*-normalized low-pass filters used for the PD-offset *τ*_b_ input. **(F)** As in (C), but for manipulations of the *τ*_b_ input. Speeding up the filter dynamics (*τ*_b_ = 75 ms) shifted model responses to higher velocities, while slowing down filter dynamics (*τ*_b_ = 225 ms) shifted responses to lower velocities. Both results are inconsistent with our experimental findings. **(G)** As in (D). **(H)** As in (E), but with the ℓ*_2_*-normalized low-pass filters of varying time constants replaced by ℓ*_2_*-normalized high-pass filters. **(I)** As in (F), but for the case in which the filter of the PD-offset input is high-pass. The responses of the ‘default’ model, in which this input has a low-pass filter with a time constant of *τ*_b_ = 150 ms, are plotted in *black*. Replacing this low-pass filter with a high-pass filter of the same time constant reverses the model’s direction preference, with responses to motion in the former ND now being greater than those to motion in the former PD. Reducing the time constant of this filter to *τ*_b_ = 75 ms exacerbates this effect. **(J)** Manipulations of the PD-offset ON inhibitory input in a synaptic model with an additional, parallel PD-offset ON inhibitory input. As in (A), only one of the two parallel inputs is manipulated at a time. **(K)** As in (H), but for the model with parallel PD-offset ON inhibitory inputs shown in (J). The filter of the non-manipulated input of this pair is kept as low-pass. **(L)** As in (I), but for the model described in (J-K). When a parallel PD-offset ON inhibitory delayed input is added, the reversal of direction preference observed in (I) no longer occurs. When the time constant of the manipulated input is equal to that of the other inputs (*τ*_b_ = 150 ms), exchanging its low-pass filter for a high-pass filter increases the model’s preferred velocity. However, when the high-pass filter’s time constant is made faster (*τ*_b_ = 75 ms), the model’s sensitivity shifts to slower velocities. The latter of these simulations is consistent with our experimental findings.

**Figure S15.**
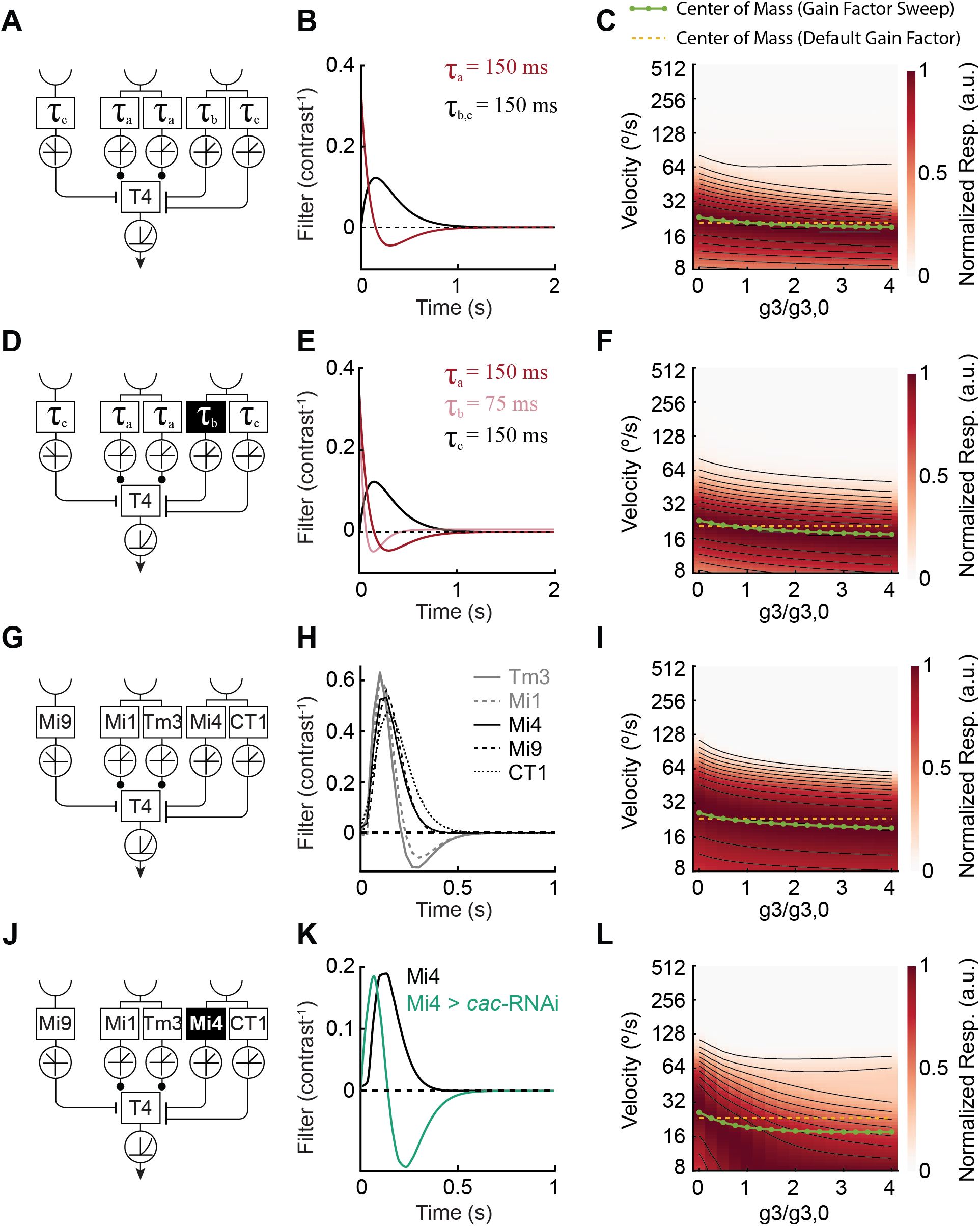
Changes in filter’s gain factor tune synaptic model responses to higher motion velocity. **(A)** Three-input synaptic model with an additional, parallel delayed PD-offset ON inhibitory input was tested with default filters and filter time constants for each input. A central, excitatory spatial input is composed of two arms, each with the same temporal dynamics (τ_a_). Two parallel PD-offset, inhibitory arms (τ_b_ and τ_c_) share one spatial receptive field. An ND-offset OFF inhibitory input has dynamics τ_c_. **(B)** Synthetic ℓ*_2_*-normalized low-pass (τ_b,c_ = 150 ms) and high-pass (τ_a_ = 150 ms) filters used in the ‘wildtype’ synaptic model. **(C)** Sweep of fractional rescaling of the τ_b_ input’s gain factor relative to its wildtype value. Tuning curves for each gain factor are shown in false color, with responses normalized by the maximal response for that gain factor. To quantify the resulting changes in tuning, the log-velocity center of mass of the wildtype model’s tuning curve (*yellow dashed line*) is compared the log-velocity centers of mass for models with altered gain factors (*green dotted line*). Here and below, decreasing the gain *g*_3_ of the PD-offset inhibitory input to T4 relative to its ‘default’ value *g*_3,0_ tended to shift T4 tuning to higher velocities. Thus, the decrease in Mi4 and CT1 filter amplitude (ignoring changes in dynamics) under *cac*-RNAi manipulation would not, according to this model, be expected to shift tuning curves to slower velocities, as observed in experiments. **(D)** As in (A), but with manipulation of the τ_b_ input filter as in Figure S14G. **(E)** As in (B), but with the filter set used in Figure S14G-L. **(F)** As in (C), but with the filters shown in (E). **(G)** As in (A), but for a model using data-driven filters (see Methods) of Mi9, Mi1, Tm3, Mi4, and CT1. **(H)** Data-driven filters used to test model described in (G). **(I)** As in (C), but using the filters of (H). Here, the gain factor for the Mi4-like input is manipulated, while that for the CT1-like input is kept fixed. **(J)** As in (A), but the Mi4-like input is manipulated with the Mi4 > *cac*-RNAi wildtype filter. **(K)** Data-driven filters of Mi4 and Mi4 > *cac*-RNAi used to test the model described in (J). **(L)** As in (I), but using the data-driven filters of (K).

